# Chemogenetic Inhibition of Lateral Hypothalamus Inputs to the Paraventricular Thalamus Reduces Goal-Tracking in a Behaviorally Flexible Subgroup of Male Rats

**DOI:** 10.64898/2026.05.28.728487

**Authors:** Amanda G. Iglesias, Jasmine K. Bhatti, Alexandra E. Turfe, Stephen Chang, Jessica Liu, Paolo Campus, Shelly B. Flagel

## Abstract

The paraventricular nucleus of the thalamus (PVT) has emerged as an important node in circuits regulating motivated behavior. The neuronal pathway from the lateral hypothalamus (LH) to the PVT has specifically been shown to regulate arousal, feeding, and reward seeking. However, the involvement of the LH–PVT pathway in individual differences in cue-motivated behavior remains unclear. During a Pavlovian conditioned approach paradigm, when a reward is repeatedly preceded by the presentation of a cue, rats come to exhibit a conditioned response to the cue. One extreme of the population, sign-trackers (STs), approach and interact with the cue itself; while the other extreme, goal-trackers (GTs), approach the location of reward delivery. Intermediate responders (IRs) approach and interact with both the cue and reward location, without a clear preference. We utilized a Pavlovian conditioned approach paradigm to examine the effects of LH–PVT pathway inhibition on individual differences in cue-motivated behavior. A dual-vector approach was used to selectively express inhibitory chemogenetic receptors in the LH–PVT pathway. We found that inhibition of the LH–PVT pathway selectively attenuates the expression of goal-tracking behavior, without affecting sign-tracking. This effect is driven primarily by IR rats, as inhibition of LH–PVT neurons attenuates goal-tracking behavior in IRs, without impacting the response of STs or GTs. We speculate that the flexibility of responding in IR rats made them especially vulnerable to this manipulation. These findings identify the LH–PVT pathway as a selective contributor to reward-directed conditioned responding and a circuit substrate for behavioral flexibility.

**Significance Statement:** Individuals differ in how reward-predictive cues motivate behavior, a feature linked to vulnerability and resilience to maladaptive reward seeking. The paraventricular thalamus (PVT), a midline hub with broad limbic connectivity, and its input from the lateral hypothalamus (LH) influences arousal, feeding, and reward seeking, but the role of the LH–PVT pathway in individual variability in cue-motivated behavior is unclear. We used chemogenetics to selectively suppress LH–PVT neurons during a Pavlovian conditioned approach paradigm. Inhibiting this projection reduced goal-tracking without altering sign-tracking, and this effect was driven primarily by intermediate responders, a phenotype characterized by behavioral flexibility. These results implicate LH–PVT signaling in reward-directed conditioned responding and suggest that this pathway contributes to flexible cue-motivated behavior.

## Introduction

Decades of work have sought to identify neural circuits that contribute to motivated behavior and reward learning, much of it focused on the classic mesocorticolimbic dopamine system (Berridge, 2007; Berridge & Robinson, 1998; Bromberg-Martin et al., 2010). Yet, motivated behavior arises from a broader network of cortical and subcortical structures. Within this network, the paraventricular nucleus of the thalamus (PVT) has emerged as an important node implicated in arousal (Hsu et al., 2014; Kirouac, 2015), reward seeking (Kirouac, 2015; Matzeu et al., 2014), stress responsivity (Bhatnagar & Dallman, 1998; Bhatnagar & Dallman, 1999; Bhatnagar et al., 2003; Bhatnagar et al., 2002; Heydendael et al., 2011; Li et al., 2010b), motivational conflict (Choi et al., 2019; Choi & McNally, 2017), and cue-motivated behavior (Beas et al., 2024; Choi et al., 2010; Igelstrom et al., 2010; Matzeu et al., 2017; Schiltz et al., 2007). Foundational work by Kelley and colleagues placed the PVT within a hypothalamic-thalamic-striatal circuit relevant for motivated behavior (Kelley et al., 2005). Anatomically, the PVT receives inputs from cortical, hypothalamic, and brainstem regions and projects to limbic targets including the nucleus accumbens (NAc), bed nucleus of the stria terminalis, and central amygdala (Hsu & Price, 2009; Li & Kirouac, 2008). This connectivity positions the PVT to integrate arousal- and state-related signals with environmental stimuli to influence motivated behavior (Barson et al., 2020; Iglesias & Flagel, 2021; Kirouac, 2015; Penzo & Gao, 2021).

Among the subcortical regions that densely innervate the PVT, the lateral hypothalamus (LH) is of particular interest (Kirouac, 2015; Kirouac et al., 2005). The LH has long been recognized for its role in appetitive and consummatory processes (Hoebel & Teitelbaum, 1962; Margules & Olds, 1962), including classic findings that LH stimulation elicits intense feeding (Anand & Brobeck, 1951; Delgado & Anand, 1953). However, the LH is now recognized as a heterogeneous structure supporting functions beyond feeding and energy balance (Bonnavion et al., 2016; Chen et al., 2025; Petrovich, 2018; Stuber & Wise, 2016). LH inputs to the PVT include orexinergic and other neurochemically diverse populations implicated in feeding, arousal, and responses to salient sensory stimuli (Kirouac et al., 2005; Otis et al., 2019; Ren et al., 2018). Yet, the contribution of the LH–PVT pathway to Pavlovian cue-motivated behavior remains unclear.

A Pavlovian conditioned approach (PavCA) paradigm can capture individual variability in conditioned responses to a reward-paired cue (Flagel et al., 2007; Hughson et al., 2019; Meyer et al., 2012; Robinson & Flagel, 2009) and has proven useful for elucidating the brain mechanisms underlying cue-motivated behavior (Campus et al., 2019; Flagel, Cameron, et al., 2011; Flagel, Clark, et al., 2011; Haight et al., 2020). In this procedure, a lever-cue (conditioned stimulus, CS) precedes the delivery of food reward (unconditioned stimulus, US), and animals develop distinct conditioned responses directed toward either the cue itself (sign-tracking) or the site of reward delivery (goal-tracking). These conditioned responses reflect differences in the value attributed to the cue (Robinson & Flagel, 2009). For sign-trackers (STs), the lever-cue acquires both predictive and incentive value, whereas for goal-trackers (GTs), it primarily acquires predictive value. The attribution of incentive motivational value, or incentive salience, transforms the cue into an attractive and desirable stimulus capable of controlling behavior in STs (Berridge et al., 2009; Flagel et al., 2009; Robinson & Berridge, 1993). Intermediate responders (IRs), a less well-studied phenotype, show greater behavioral flexibility, vacillating between sign- and goal-tracking behavior.

Prior work has identified the PVT as an important node in the regulation of sign- and goal-tracking behaviors (Campus et al., 2019; Flagel, Cameron, et al., 2011; Haight & Flagel, 2014; Haight et al., 2015; Haight et al., 2017; Kuhn et al., 2018). A top-down cortico-thalamic pathway has been proposed to encode the predictive value of reward cues and regulate conditioned responding across phenotypes, whereas bottom-up inputs to the PVT are thought to relay the incentive value of reward-associated cues (Campus et al., 2019; Haight & Flagel, 2014; Haight et al., 2017). Within this framework, the LH is of particular interest, as it sends dense orexinergic projections to the PVT (Kirouac et al., 2005; Peyron et al., 1998). Consistent with this idea, LH lesions attenuate the development of sign-tracking behavior, and pharmacologically blocking orexin signaling in the PVT reduces sign-tracking and the incentive motivational value of the lever-cue (Haight et al., 2020).

Here, we examined the role of the LH–PVT pathway in the expression of cue-motivated behavior in male rats using a chemogenetic approach. We assessed the effects of LH–PVT inhibition across the population and then examined whether those effects varied by PavCA phenotype. Based on prior work, we hypothesized that inhibiting LH–PVT projections would reduce sign-tracking by diminishing the incentive value of the cue. Instead, LH–PVT inhibition selectively attenuated goal-tracking, and this effect was driven by IRs, with no detectable effect within ST or GT populations.

## Materials and Methods

Behavioral testing took place during the light-phase between 10 AM and 4 PM. All procedures followed The Guide for the Care and Use of Laboratory Rats: Eighth Edition (2011, National Academy of Sciences) and were approved by the University of Michigan Institutional Animal Care and Use Committee. Rats were run in two separate experiments. Experiment 1 assessed the effect of chemogenetic inhibition of the LH–PVT pathway on PavCA behavior in rats expressing inhibitory DREADD (Designer Receptors Exclusively Activated by Designer Drugs) receptors. Experiment 2 served as a no-DREADD control experiment used to determine whether clozapine-N-oxide (CNO; DREADD ligand) in the absence of DREADD affected PavCA behavior. In Experiment 1, rats underwent surgery to express DREADD receptors in the LH–PVT pathway and were run across multiple rounds, with either 30 or 40 rats tested per round. In Experiment 2, rats did not undergo surgery or express DREADD receptors and were run in one round of 60 rats.

The behavioral timeline was the same for both experiments (Figure 1). Rats underwent 7 consecutive sessions of PavCA. We aimed to test the effects of LH–PVT pathway inhibition during the expression of PavCA behavior. After sessions 3–4 of PavCA (acquisition) rats were given vehicle (VEH) injections to acclimate them to receiving injections. Twenty-five minutes prior to sessions 5–7 of PavCA (expression), rats were either given VEH or CNO. On session 8 all rats underwent a conditioned reinforcement test (CRT), with no injections given prior to testing.

**Figure 1.**
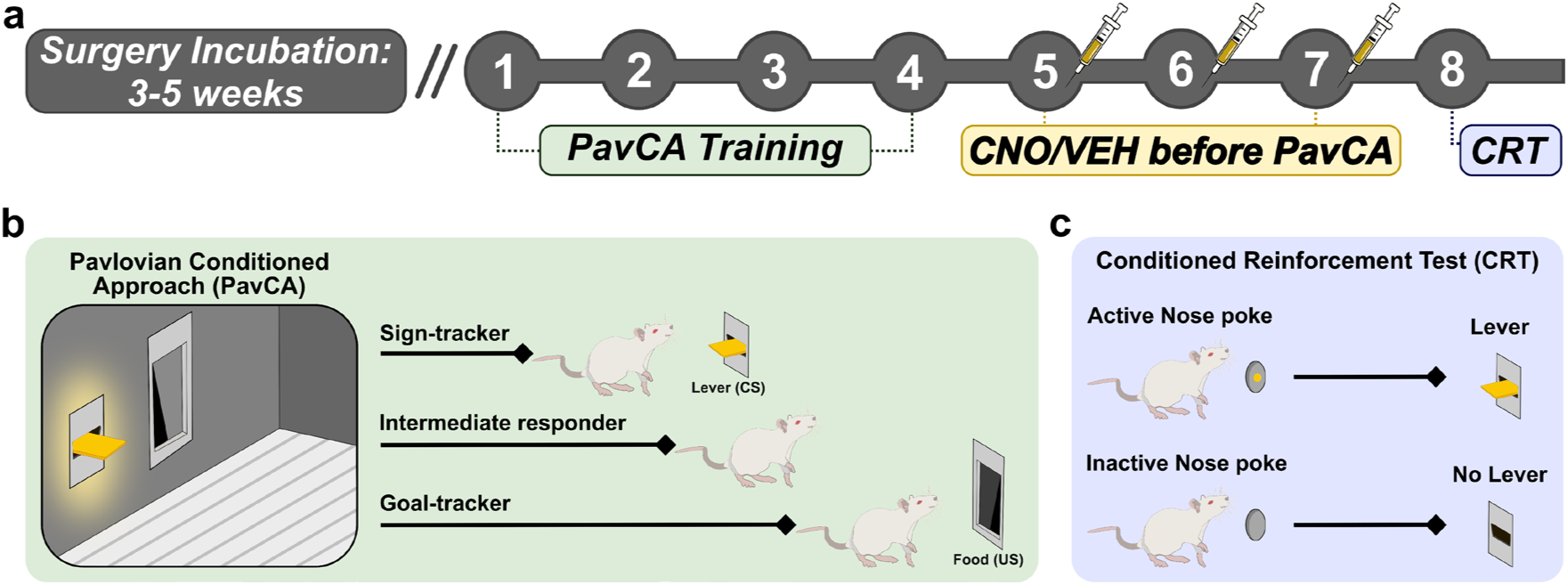
Experimental timeline. (a) Schematic of the experimental timeline: Stereotaxic surgery was performed to infuse inhibitory DREADD (Designer Receptors Exclusively Activated by Designer Drugs) virus in the lateral hypothalamus (LH) to the paraventricular nucleus of the thalamus (PVT) pathway. The virus incubated in the brain for 3–5 weeks and was followed by behavioral assays. Pavlovian conditioned approach (PavCA) training occurred for 7 consecutive days. On days 5–7 of behavior, either clozapine-N-oxide (CNO; 5 mg/kg) or vehicle (VEH; 6% dimethyl sulfoxide) was injected intraperitoneally. On day 8 of behavior rats had a conditioned reinforcement test (CRT) with no injections immediately prior. (b) PavCA Schematic illustrating the behavior box and responses of sign-tracker, intermediate responder, and goal-tracker. (c) CRT schematic wherein nose pokes in the port designated active result in lever presentation and those in the inactive port are without consequence.

### Subjects

Rats were approximately 9 weeks of age (∼300–350 g body weight) at the time of arrival and were housed under a 12-hour light-dark cycle (lights on at 6 or 7 AM depending on daylight saving time) with climate-controlled conditions (22 ± 2°C). They had *ad-libitum* access to food and water throughout the study. Rats were pair-housed for the duration of the experiment and were acclimated to the housing room for a minimum of one week before experimenter handling began.

Male Sprague Dawley rats (N = 230 in total) were ordered from different barriers at either Charles River Laboratories or Taconic Biosciences. In Experiment 1 (5 rounds, n = 170), rats were from Charles River barrier R04 (n = 41, Raleigh, NC), Charles River barrier R08 (n = 64, Raleigh, NC), Taconic BU16 (n = 20, Cambridge City, IN), or Taconic IBU (n = 45, Germantown, NY). Seven rats were removed from the study before behavioral testing due to surgical or post-surgical complications. Rats were characterized as STs, IRs, or GTs, based on PavCA behavior on session 4, prior to any CNO administration. Rats were then assigned to VEH or CNO treatment groups in a counterbalanced manner within each phenotype. To avoid overrepresentation of a single phenotype within a given round, rats (n = 66) from overrepresented phenotypes were randomly excluded prior to DREADD activation and before any test-phase behavior was analyzed. Additonally, rats (n = 43) were excluded for off-target viral expression or minimal expression in the LH, aPVT, pPVT, or bilaterally across the targeted pathway (see Immunohistochemistry). These behavioral and anatomical exclusions were applied in sequence and contributed to the final sample sizes. A total of n = 54 rats were included in Experiment 1, as follows: STs (n = 13; 7 VEH/6 CNO), IRs (n = 18; 7 VEH/11 CNO), or GTs (n = 23; 15 VEH/8 CNO). In Experiment 2, the no-DREADD cohort (n = 60), rats were either from Charles River barrier R04 (n = 30, Raleigh, NC) or Charles River barrier R08 (n = 30, Raleigh, NC). We ordered from Charles River barriers for this cohort because, historically, these barriers yielded a more even distribution across phenotypes. A total of n = 60 rats were used for Experiment 2, as follows: STs (n = 21; 11 VEH/10 CNO), IRs (n = 23; 11 VEH/12 CNO), or GTs (n = 16; 8 VEH/8 CNO).

### Viral Vectors

Viruses were obtained from Addgene. The Cre-dependent G_i_ DREADD virus (AAV8-hSyn-DIO-hM4Di-mCherry, Addgene 44362-AAV8) had a titer of ≥ 1 × 10^13^ vg/mL. The Cre virus (AAVrg-pmSyn1-EBFP-Cre, Addgene 51507-AAVrg) had a titer of ≥ 7 × 10^12^ vg/mL.

### DREADD ligand

We obtained CNO from the National Institute of Mental Health. DREADD receptor activation occurred via systemic administration of CNO, which results in inhibition of the LH–PVT pathway through disruption of G-protein-dependent signaling, likely from induced hyperpolarization (Armbruster et al., 2007; Vardy et al., 2015) or inhibition of presynaptic release of neurotransmitters (Stachniak et al., 2014; Vardy et al., 2015). CNO (5 mg/kg) was dissolved in 6% dimethyl sulfoxide (DMSO) and sterile H_2_O and injected intraperitoneally (i.p.) (Armbruster et al., 2007; Ferguson et al., 2011; Rogan & Roth, 2011; Stachniak et al., 2014). Control animals were administered vehicle i.p. (VEH; 6% DMSO and sterile H_2_O).

### Surgery

Surgeries were performed using a dual-vector approach to selectively express hM4Di-DREADD receptors (G_i_, inhibitory) in the LH–PVT pathway. Rats were anesthetized with 5% isoflurane (Vet One, Boise, ID) delivered via an induction chamber and were then placed into the stereotaxic frame (David Kopf instruments, Tujunga, CA or Stoelting, Wood Dale, IL) and given an injection of carprofen (5 mg/kg, subcutaneous (s.c.)) for analgesia. Rats remained at 2% isoflurane for anesthetic maintence. Surgeries were performed under aseptic conditions. All coordinates were relative to bregma (Paxinos & Watson, 2007). A Neuros Syringe (5 µL Model 33-gauge, point style 3, Hamilton Company) infused 0.75 µl of an inhibitory Cre-dependent Gi DREADD virus bilaterally into the LH (−2.7 mm AP (anterior/posterior); ±1.6 mm ML (medial/lateral); −9 mm DV (dorsal/ventral)). The DREADD virus was infused over 8 min and the syringe was left in place for 10 additional min. A Neuros Syringe (0.5 µL Model 33-gauge, point style 3, Hamilton Company) infused 0.10 µl of a retrograde Cre virus at a 10° angle into the anterior PVT (aPVT: −2.0 mm AP; −1 mm ML; −5.4 mm DV) and posterior PVT (pPVT: −3.0 mm AP; −1 mm ML; −5.5 mm DV) to target the entire rostro-caudal axis of the region. The Cre virus was infused over 2 min and the syringe was left in place for 5 additional min. Following 3–5 weeks of virus incubation, all rats underwent Pavlovian conditioned approach training.

### Pavlovian conditioned approach (PavCA)

Pavlovian conditioned approach (PavCA) training occurred inside Med Associates chambers (St. Albans, VT, USA; 24.1 × 21 × 30.5 cm) located in sound-attenuating boxes equipped with a fan to reduce background noise. The chambers contained a food cup that was connected to a pellet dispenser and placed in the center of one wall 3 cm above the chamber floor. An illuminated retractable lever-cue was located either to the left or right of the food cup, 6 cm above the chamber floor. A white house light was placed at the top of the chamber on the wall opposite the food cup and lever-cue and remained on for the duration of the session. Food cup entries were recorded upon a photo-beam break inside the food cup, and a lever-cue contact was recorded following a minimum deflection force of 10 g.

All rats were handled by experimenters for two days before assessing PavCA behavior. For the two days prior to pre-training, rats received ∼25 banana-flavored grain pellets (each pellet 45-mg; Bio-Serv, Flemington, NJ, USA) in their home cage to familiarize them with the reward. Prior to each session, the rats were transferred to the injection room 1 hour prior to behavior. Rats were initially placed into the Med Associates chambers for a single pretraining session. At the start of the pretraining session, the food cup was baited with two food pellets to direct the rats’ attention to the location of reward delivery. During pretraining, the lever-cue remained retracted, and rats received a food pellet in the food cup on a variable 30 s (range 0–60 s) schedule. There were 25 trials, and the pretraining session lasted approximately 12.5 min, during which food cup entries were recorded, and food pellet consumption was confirmed. Following pretraining, rats had a single session of PavCA each day for 7 consecutive days (Figure 1a,b). The acquisition phase of PavCA behavior occurred across the first 4 sessions. Following behavior on sessions 3–4, rats received i.p. VEH injections to acclimate them to receiving injections. Rats were assigned to phenotype and treatment groups in a counterbalanced manner based on their behavior on session 4. On sessions 5–7, rats received i.p. CNO injections to activate the DREADDs 25 min before the start of the session each day. Each PavCA session began with a 5-min waiting period, followed by illumination of the house light to signify the start of the session. During PavCA, the illuminated lever-cue (CS) entered the chamber for 8 s, and upon retraction, a food pellet (US) was immediately delivered into the adjacent food cup. PavCA sessions consisted of 25 lever-cue (CS)/food-US trials on a variable 90 s schedule (range 30–150 s) (Campus et al., 2019; Meyer et al., 2012). Each session lasted approximately 40 min. It was confirmed that all food pellets had been consumed following each session.

Med Associates software recorded the following behaviors during PavCA sessions: (1) number of food cup entries made during the 8-s lever presentation, (2) latency to contact the food cup during lever-cue presentation, (3) number of lever-cue contacts, (4) latency to lever-cue contact, and (5) the number of food cup entries made during the inter-trial interval (i.e., food cup entries made in between lever-cue presentations). Contact, latency, and probability data were used to calculate the daily PavCA score, and the score from session 4 alone was used to characterize rats into their behavioral phenotypes (Meyer et al., 2012). The PavCA score is a composite measure calculated using the following formula: [(Probability Difference Score + Response Bias Score + (-Latency Difference Score))/3]. PavCA score values range from −1 to 1, with −1 representing individuals with a conditioned response (CR) focused solely on the food cup during lever-cue presentation (i.e., a “pure” goal-tracker, GT) and +1 representing individuals with a CR focused solely on the lever-cue upon its presentation (i.e., a “pure” sign-tracker, ST). Intermediate responders (IRs) vacillate between the two responses. The PavCA Index was generated from the behavioral measures during session 4 to classify animals as STs (PavCA ≥ 0.5), IRs (−0.5 < PavCA < 0.5), or GTs (PavCA ≤ −0.5).

### Conditioned reinforcement test (CRT)

Rats were exposed to a conditioned reinforcement test (CRT) the day following the final PavCA session (Figure 1a,c). This test required rats to perform an instrumental response for lever presentation, in the absence of reward (Robinson & Flagel, 2009). That is, the lever that was previously the reward cue now served as the reinforcer. Conditioning chambers were reconfigured, and the lever was placed in the center wall, flanked by nose ports. Nose pokes into the “active” port resulted in presentation of the illuminated lever for 2 s, while nose pokes into the “inactive” port had no consequence. The active port was placed opposite the side of the lever location during PavCA sessions to minimize side bias. The conditioned reinforcement test lasted 40 min. Med Associates software recorded the number of nose pokes into the “active” and “inactive” ports and the number of contacts with the lever during its brief presentation.

### Perfusion and tissue processing

Rats that previously received surgery were perfused within 5 days following the experiment. Rats were anesthetized with a cocktail of ketamine (90 mg/kg, i.p.) and xylazine (10 mg/kg, i.p.). Ten min after the anesthetic injection, rats were transcardially perfused with 0.9% saline and 4% formaldehyde (pH = 7.4). Following brain extraction, the tissue was post-fixed in 4% formaldehyde for 24 hr at 4°C and then placed in 30% sucrose at 4°C (sucrose in 0.1M PBS, pH = 7.4) for 3 days. The brains were frozen using dry ice and coated in Tissue-Plus Optimal Cutting Temperature (OTC) compound (Fisher HealthCare, Houston, TX). Serial coronal brain slices were taken at 40 µm using a cryostat at −20°C (Leica Biosystems Inc, Buffalo Grove, IL).

### Immunohistochemistry

DREADD expression in LH–PVT projection neurons was evaluated following immunohistochemical staining of mCherry, the fluorescent tag on the G_i_ DREADD virus. Free-floating coronal sections from the LH and PVT were initially washed 6 times in 0.1M PBS (pH = 7.4) to remove Optimal Cutting Temperature (OTC) debris. Each step after this initial wash was followed by 3 washes in 0.1M PBS (pH = 7.4). Sections were incubated in 1% H_2_O_2_ for 10 min to block endogenous peroxidase activity and then incubated in 0.1M PBS containing 0.4% Triton X-100 (TX) and 2.5% Normal Donkey Serum (NDS) (Jackson ImmunoResearch Laboratories, Inc, West Grove, PA) to block nonspecific binding of the secondary antibody. Subsequently, sections were incubated overnight at room temperature in primary antibody (rabbit anti-mCherry, Abcam, Cambridge, UK, diluted 1:30,000) in 0.1M PBS + 0.4% TX + 1% NDS, and the next day sections were incubated for 1 hr in a biotinylated donkey anti-rabbit secondary antibody (Jackson Immunoresearch, West Grove, PA, diluted 1:500) in 0.1M PBS + 0.4% TX + 1% NDS. Sections were then incubated for 1 hr in Vectastain Elite ABC solution (1:1000 A and 1:1000 B, diluted in 0.1M PBS, mixed 30 min before use; Vector Laboratories). This stain was visualized by incubating the sections in 0.1M NaPB containing 0.02% 3,3’-diaminobenzidine tetrahydrochloride (Sigma-Aldrich) and 0.012% H_2_O_2_ for 10 min causing a brown precipitate to form at the location of mCherry detection. Sections were stored at 4°C until mounted, air dried, and cover slipped with Permount (Thermo-Fisher Scientific, Waltham, MA). Bright-field images containing the LH, aPVT, and pPVT were captured using a Leica DM1000 light microscope (Leica-Microsystems, Wetzlar, GER) and were analyzed by an experimenter blind to the experimental groups.

The experimenter assigned a score of 0–3 to each image according to both the density and location of DREADD expression in the areas of interest (Figure 2). A score of 0 was assigned to subjects that had no DREADD expression in the LH, aPVT, and pPVT or off-target DREADD expression (e.g. expression outside regional boundaries); a score of 1 was assigned to subjects that had visible DREADD expression in both the LH and PVT; a score of 2 was assigned to subjects with visible DREADD expression in one region and strong DREADD expression in the other; a score of 3 was assigned to subjects that had strong DREADD expression in both the LH and PVT. Rats that had a score of 0 (n = 43), meaning they had no visible expression in either the LH or PVT, were excluded from the statistical analysis. Because the LH and zona incerta (ZI) are anatomically adjacent, some mCherry-positive cell body expression was observed in the ZI in included subjects. This expression was unavoidable given the proximity of the targeted LH injection site to the ZI. Rats were excluded only when viral expression was absent from the LH/PVT pathway or when expression was judged to be off-target relative to the intended LH–PVT targeting strategy.

**Figure 2.**
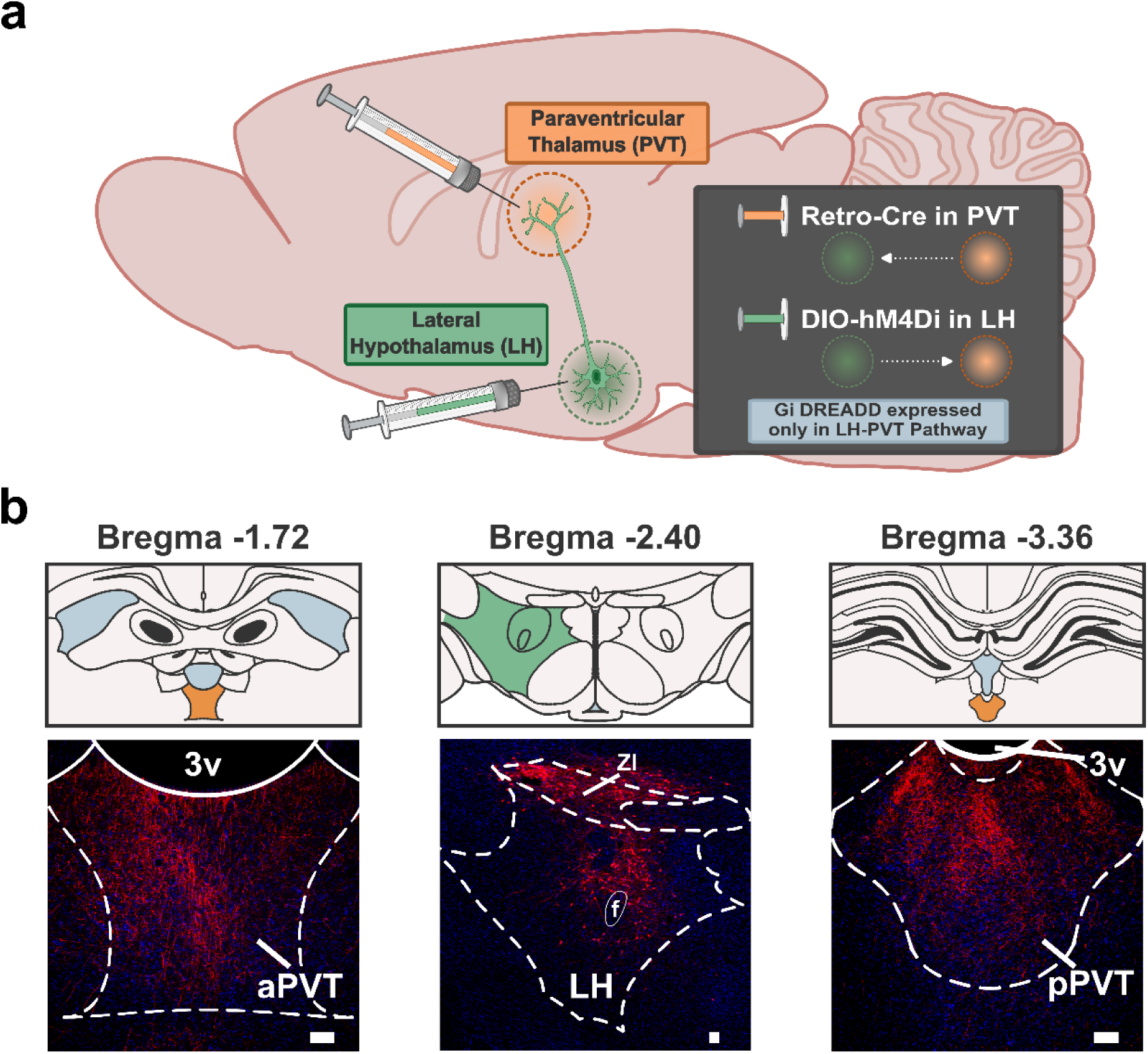
Surgery schematic and virus expression. (a) Schematic of the dual-vector strategy used to selectively express inhibitory DREADD in the lateral hypothalamus-paraventricular nucleus of the thalamus (LH–PVT) pathway. A retrograde Cre virus was infused into the anterior and posterior PVT (aPVT and pPVT), and a Cre-dependent inhibitory DREADD virus was infused bilaterally into the LH. (b) Representative images showing DREADD virus expression in the aPVT (10× magnification), LH (4× magnification), and pPVT (10× magnification). Scale bars in the bottom right corner represent 100 µm. 3v, third ventricle; ZI, zona incerta; f, fornix.

To obtain representative images and visualize mCherry expression, a subset of brains were processed for immunofluorescence. Free-floating coronal sections from the LH and PVT were initially washed 6 times in 0.1M PBS (pH = 7.4) to remove OTC debris. Each step after this initial wash was followed by 3 washes in 0.1M PBS (pH = 7.4). Sections were incubated overnight at room temperature in primary antibody (rabbit anti-mCherry, Abcam, Cambridge, UK, diluted 1:500) in 0.1M PBS + 0.4% TX + 2.5% NDS. The next day sections were incubated for 2 hr in a biotinylated donkey anti-rabbit secondary antibody (Jackson Immunoresearch, West Grove, PA, diluted 1:500) in 0.1M PBS + 0.4% TX + 2.5% NDS. Sections were then incubated for 1 hr in Alexa Fluor 594-conjugated streptavidin (Thermo Fisher Scientific, Waltham, MA, diluted 1:1000) in 0.1M PBS + 0.4% TX, then mounted onto slides and cover slipped with ProLong Gold Antifade Mountant (Thermo Fisher Scientific, Waltham, MA). Fluorescent images of the LH, aPVT, and pPVT were captured using a FV3000 confocal microscope and the FV31S-SW Viewer software (OLYMPUS Microscopes, Center Valley, PA, USA) (Figure 2b). ***Statistical analyses*** Statistical analyses were conducted with the Statistical Package for the Social Sciences (SPSS) program version 27.0 (IBM, Armonk, NY, USA). To assess PavCA behavioral outcome measures across sessions, a linear mixed-effects model (LMM) with restricted maximum likelihood estimation was used to account for repeated measures. This analysis applied multiple covariance structures to the data set, and the structure with the lowest Akaike information criterion (AIC) was selected as best fit (Duricki et al., 2016; Verbeke, 1997). When two variables were compared at a single timepoint or across two timepoints, ANOVAs were performed, as described below. For all analyses, statistical significance was set at *p* < 0.05, and Bonferroni *post hoc* comparisons were made when significant main effects or interactions were detected. Effect size (Cohen’s d, Cohen, 1988) was calculated for pairwise comparisons. Effect sizes were considered with respect to the following indices: 0.2, small; between 0.5 and 0.8, medium; between 1.2 and 2.0, large (Cohen, 1988; Sawilowsky, 2009).

A LMM was conducted to compare treatment (VEH vs. CNO) and phenotype (ST vs. IR vs. GT) across PavCA sessions, with sessions 1–4 or 5–7 used as the repeated variable. To ensure that subjects were counterbalanced between experimental groups after PavCA training, the average PavCA Index from session 4 was analyzed using a two-way ANOVA with treatment (VEH vs. CNO) and phenotype (ST vs. IR vs. GT) as independent variables. A two-way repeated measures ANOVA was conducted when session 4 (“pretest”) was directly compared to the average of sessions 5–7 (“test”), with session (“pretest” vs. “test”) as the within-subjects independent variable and treatment (VEH vs. CNO) and phenotype (ST vs. IR vs. GT) as the between-subjects independent variables. For both the LMM and ANOVAs, PavCA Index, inter-trial interval (ITI) food cup entries, sign-tracking behaviors (number of lever-cue contacts, probability to approach the lever-cue, latency to approach the lever-cue), and goal-tracking behaviors (food cup entries during lever-cue presentation, probability to approach the food cup during lever-cue presentation, latency to approach the food cup during lever-cue presentation) were used as dependent variables.

A mixed ANOVA was used to assess the effects of treatment (VEH vs. CNO), port (active vs. inactive), and phenotype (ST vs. IR vs. GT) on nose pokes during the conditioned reinforcement test; and an ANOVA was used to assess the effects of treatment (VEH vs. CNO) and phenotype (ST vs. IR vs. GT) on lever contacts.

## Results

### Acquisition of PavCA Behavior (Pretest)

The acquisition phase for Pavlovian conditioned approach (PavCA) occurred across sessions 1–4. The PavCA Index on session 4 was used to assign phenotypes and counterbalance treatment groups. There were no significant differences in the PavCA Score between treatment groups during sessions 1–4, nor in the PavCA Index on session 4 (Extended Data Table 1; Extended Data Figure 1a, b). The PavCA Score changed across sessions 1–4 (effect of session: *F*_3,73.871_ = 3.164, *p* = 0.029; Extended Data Figure 1a), but the lack of treatment effect indicated that the groups were equally balanced prior to DREADD activation. When the sign- and goal-tracking metrics that comprise the composite PavCA Index were assessed separately (Figure 3), treatment groups learned the cue-reward relationship at similar rates across sessions 1–4, as there was a significant effect of session for all measures, but no effect of treatment (Table 1a and 1d; Figure 3a, c, e, g, i, k). *Post hoc* comparisons revealed that engagement with the lever-cue and food cup significantly increased from session 1 to session 4 for all measures (*p* < 0.001). Thus, rats assigned to VEH and CNO groups acquired PavCA behavior similarly before any chemogenetic manipulation occurred.

**Figure 3.**
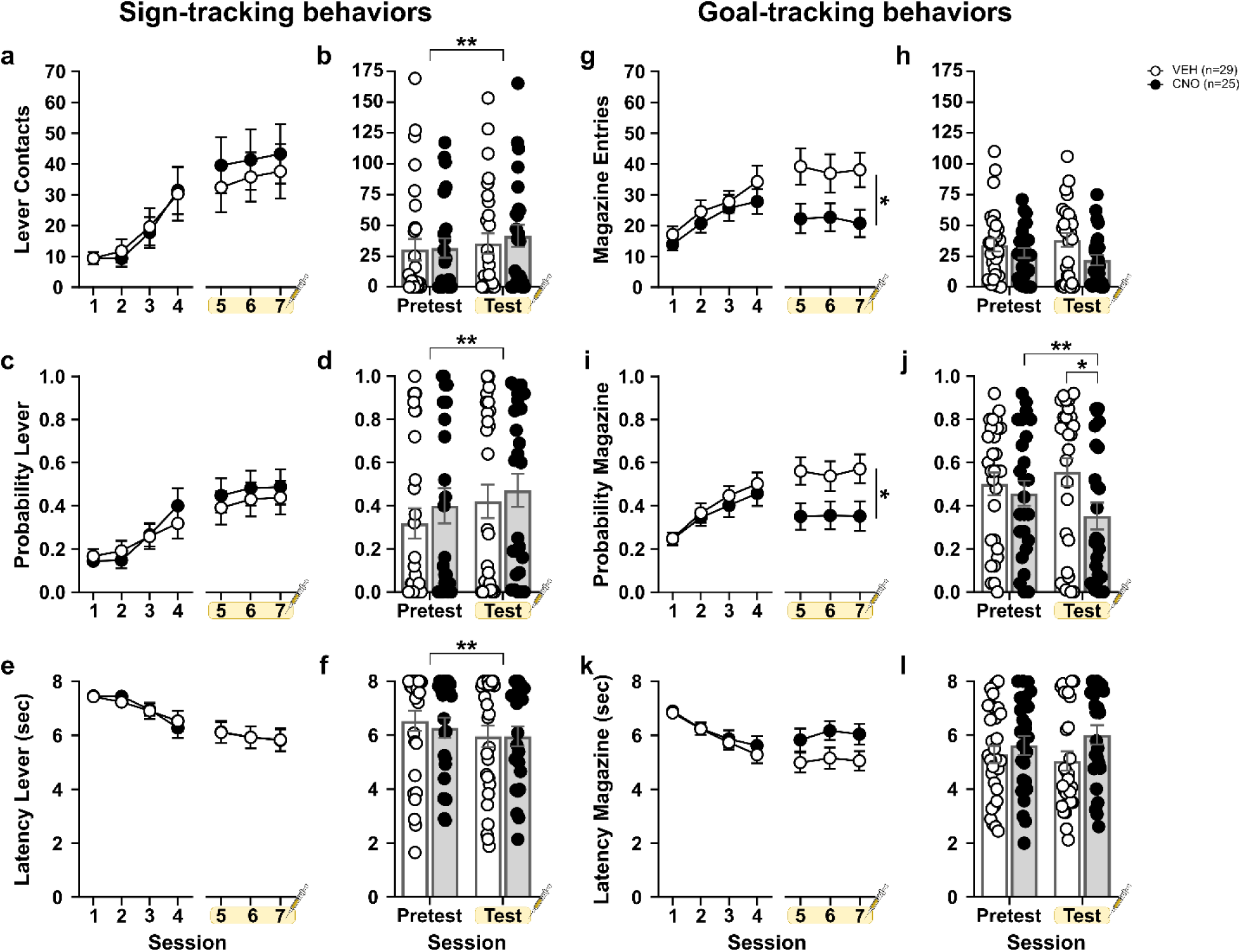
Chemogenetic inhibition of the LH–PVT pathway does not affect sign-tracking behavior, but attenuates goal-tracking behavior. (a–f) Sign-tracking and (g–l) goal-tracking behaviors expressed as mean ± SEM for (top row) number of contacts/entries, (middle row) probability to approach, and (bottom row) latency to approach the lever-cue or food cup shown across PavCA sessions 1–7 (columns 1 and 3) and as pretest (session 4) versus test (average of sessions 5–7) sessions (columns 2 and 4). On sessions 5–7, rats received either VEH or CNO prior to testing. (a–f) There were no treatment effects on sign-tracking behavior. When treatment groups were collapsed, (b) rats made more lever-cue contacts, (d) showed a greater probability of approaching the lever-cue, (f) and had decreased latency to approach the lever-cue during test sessions relative to pretest. (g, h) In contrast, CNO-treated rats showed reduced goal-tracking behavior, reflected by (g) fewer food cup entries and (i) a lower probability of approaching the food cup relative to VEH-treated rats. Graphs in b, d, f, h, j, and l include individual data points for all rats. Brackets indicate significant differences between or within pretest and test sessions; *p < 0.05, **p < 0.01.

**Table 1.** Statistical analyses for effect of treatment on sign- and goal-tracking behaviors. Data from linear mixed effects model analyses for (a,d) sessions 1–4 and (b,e) sessions 5–7, or a two-way repeated measures ANOVA (c,f) pretest vs test sessions with comparisons by treatment, collapsed across phenotype. Contacts, probability, and latency are represented for (a–c) sign-tracking behaviors and (d–f) goal-tracking behaviors. Significant effects and interactions are bolded. These data correspond to that shown in Figure 3.

|  | **a. Sessions 1–4: Sign-tracking behaviors** |  |  |  |  |  |  |  |  |
| --- | --- | --- | --- | --- | --- | --- | --- | --- | --- |
|  | Lever Contacts |  |  | Probability Lever |  |  | Latency Lever |  |  |
|  | DF | *F* | *p* | DF | *F* | *p* | DF | *F* | *p* |
| *Session* | 3,62.664 | 7.647 | **<.001** | 3,68.418 | 7.703 | **<.001** | 3,62.467 | 8.544 | **<.001** |
| *Treatment* | 1,52.222 | 0.014 | .907 | 1,52.491 | 0.008 | .929 | 1,52.285 | 0.000 | .984 |
| *Session*×*Treatment* | 3,62.664 | 0.358 | .784 | 3,68.418 | 0.972 | .411 | 3,62.467 | 1.093 | .359 |
|  | **b. Sessions 5–7: Sign-tracking behaviors** |  |  |  |  |  |  |  |  |
|  | Lever Contacts |  |  | Probability Lever |  |  | Latency Lever |  |  |
|  | DF | *F* | *p* | DF | *F* | *p* | DF | *F* | *p* |
| *Session* | 2,95.247 | 0.868 | .423 | 2,59.794 | 1.634 | .204 | 2,93.324 | 1.145 | .323 |
| *Treatment* | 1,52.028 | 0.252 | .618 | 1,52.570 | 0.228 | .635 | 1,52.066 | 0.000 | 1.000 |
| *Session*×*Treatment* | 2,95.247 | 0.038 | .963 | 59.794 | 0.009 | .991 | 2,93.324 | 0.010 | .990 |
|  | **c. Pretest vs Test: Sign-tracking behaviors** |  |  |  |  |  |  |  |  |
|  | Lever Contacts |  |  | Probability Lever |  |  | Latency Lever |  |  |
|  | DF | *F* | *p* | DF | *F* | *p* | DF | *F* | *p* |
| *Session* | 1,52 | 7.506 | **.008** | 1,52 | 8.604 | **.005** | 1,52 | 10.417 | **.002** |
| *Treatment* | 1,52 | 0.098 | .756 | 1,52 | 0.401 | .530 | 1,52 | 0.062 | .805 |
| *Session*×*Treatment* | 1,52 | 0.847 | .362 | 1,52 | 0.260 | .612 | 1,52 | 0.839 | .364 |
|  | **d. Sessions 1–4: Goal-tracking behaviors** |  |  |  |  |  |  |  |  |
|  | Food Cup Entries |  |  | Probability Food Cup |  |  | Latency Food Cup |  |  |
|  | DF | *F* | *p* | DF | *F* | *p* | DF | *F* | *p* |
| *Session* | 3,55.853 | 7.157 | **<.001** | 3,53.899 | 11.957 | **<.001** | 3,68.593 | 15.438 | **<.001** |
| *Treatment* | 1,57.616 | 0.996 | .322 | 1,58.158 | 0.347 | .558 | 1,56.378 | 0.217 | .643 |
| *Session*×*Treatment* | 3,55.853 | 0.246 | .864 | 3,53.899 | 0.132 | .941 | 3,68.593 | 0.171 | .916 |
|  | **e. Sessions 5–7: Goal-tracking behaviors** |  |  |  |  |  |  |  |  |
|  | Food Cup Entries |  |  | Probability Food Cup |  |  | Latency Food Cup |  |  |
|  | DF | *F* | *p* | DF | *F* | *p* | DF | *F* | *p* |
| *Session* | 2,69.444 | 0.159 | .853 | 2,103.058 | 0.225 | .799 | 2,51.585 | 0.928 | .402 |
| *Treatment* | 1,52.137 | 5.147 | **.027** | 1,52.025 | 5.119 | **.028** | 1,52.012 | 3.655 | .061 |
| *Session*×*Treatment* | 2,69.444 | 0.253 | .777 | 2,103.058 | 0.362 | .697 | 2,51.585 | 0.216 | .807 |
|  | **f. Pretest vs Test: Goal-tracking behaviors** |  |  |  |  |  |  |  |  |
|  | Food Cup Entries |  |  | Probability Food Cup |  |  | Latency Food Cup |  |  |
|  | DF | *F* | *p* | DF | *F* | *p* | DF | *F* | *p* |
| *Session* | 1,52 | 0.126 | .724 | 1,52 | 1.044 | .312 | 1,52 | 0.249 | .620 |
| *Treatment* | 1,52 | 3.072 | .086 | 1,52 | 2.363 | .130 | 1,52 | 1.970 | .166 |
| *Session*×*Treatment* | 1,52 | 3.487 | .067 | 1,52 | 10.532 | **.002** | 1,52 | 3.857 | .055 |

When phenotype was incorporated into the analysis for PavCA Score, there were no significant effects of treatment, but there were significant differences between phenotypes, as expected (effect of session: *F*_3,65.959_ = 3.614, *p* = 0.018; effect of phenotype: *F*_2,50.528_ = 42.631, *p* < 0.001; session × phenotype interaction: *F*_6,65.959_ = 20.242, *p* < 0.001; Extended Data Figure 1c). In agreement, the treatment groups within each respective phenotype had similar rates of sign-tracking (Table 2a; Figure 4a, c, e) and goal-tracking (Table 3a; Figure 4g, i, k) behaviors, evidenced by significant effects of session and phenotype as well as a significant session × phenotype interaction for all measures, but no effect of treatment (Table 2a, 3a). *Post hoc* comparisons revealed that, by session 4, STs, IRs, and GTs were all significantly different from one another across all sign-tracking (*p* < 0.001) and goal-tracking measures (*p* < 0.001) and on the composite PavCA Index (*p* < 0.001). However, phenotypes did not differ by treatment on session 4 (e.g., VEH STs were similar to CNO STs, etc.) (Extended Data Figure 1d). Thus, treatment groups were adequately counterbalanced within phenotype, and any changes in behavior between treatment groups thereafter would presumably be due to LH–PVT inhibition.

**Figure 4.**
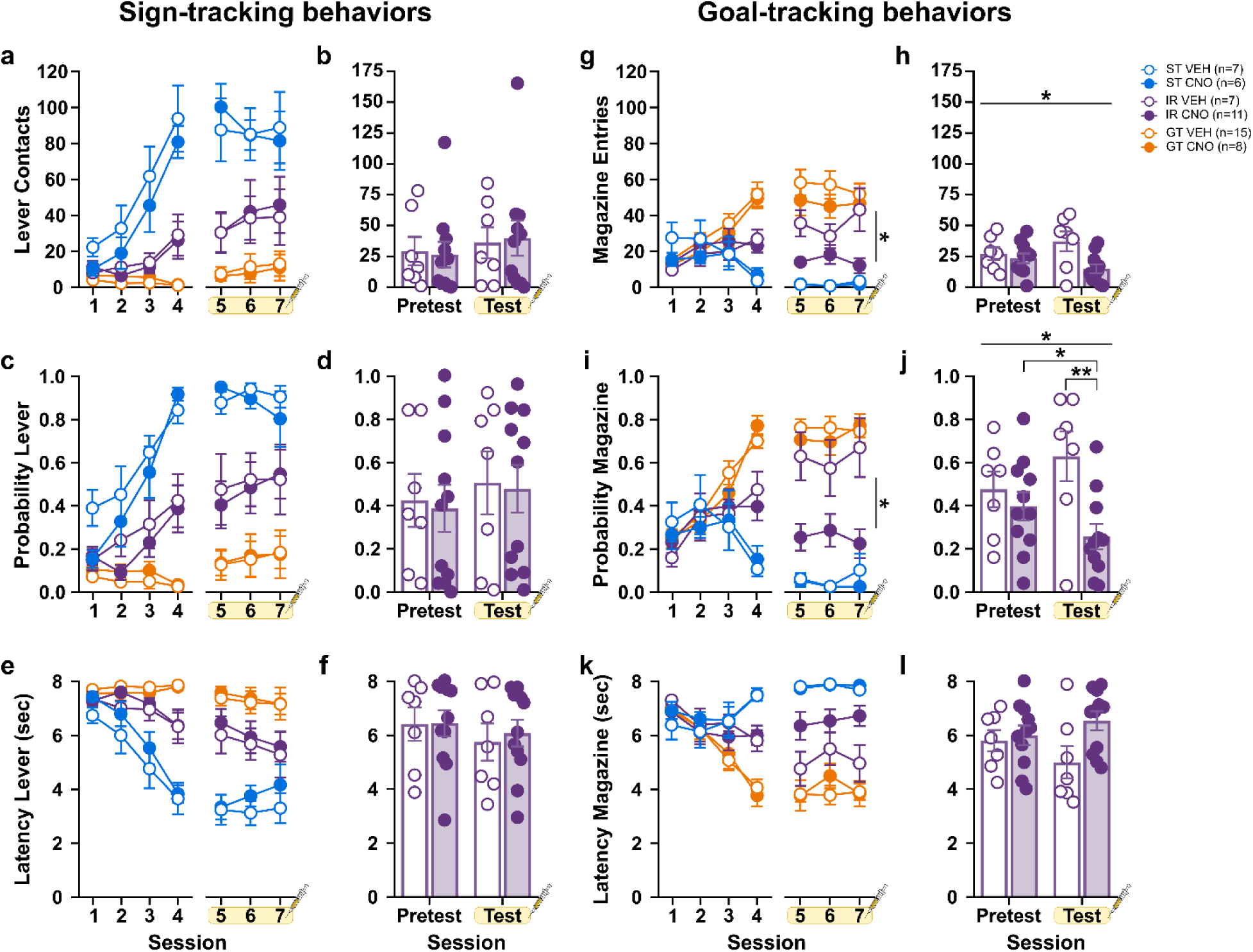
Chemogenetic inhibition of the LH–PVT pathway attenuates goal-tracking behavior in intermediate responders. (a–f) Sign-tracking behaviors and (g–l) goal-tracking behaviors are expressed as mean ± SEM for (top row) number of contacts/entries, (middle row) probability to approach, and (bottom row) latency to approach the lever-cue or food cup across PavCA sessions 1–7 (columns 1 and 3) and as pretest (session 4) versus test (average of sessions 5–7) sessions (columns 2 and 4). On sessions 5–7, rats received either VEH or CNO prior to testing. (a–f) There were no treatment effects on sign-tracking behavior within any of the phenotypes. (g–h) In contrast, CNO-treated intermediate responders (IRs) showed attenuated goal-tracking behavior, reflected by (g) fewer food cup entries and (j) a lower probability of approaching the food cup relative to VEH-treated IRs. No treatment effects were observed in sign-trackers (STs) or goal-trackers (GTs) on goal-tracking indices. Graphs in b, d, f, h, j, and l show pretest versus test comparisons and include individual data points for IR rats. Straight lines indicate a significant main effect of treatment. Brackets indicate significant differences between or within pretest and test sessions; *p < 0.05, **p < 0.01.

**Table 2.** Statistical analyses for effects of treatment and phenotype on sign-tracking behaviors. Data from linear mixed effects model analyses for (a) sessions 1–4 and (b) sessions 5–7, with comparisons by treatment and phenotype. A two-way repeated measures ANOVA assessing the effect of treatment within intermediate responders during (c) pretest vs test sessions. Contacts, probability, and latency are represented for sign-tracking behaviors. Significant effects and interactions are bolded. These data correspond to that shown in Figure 4.

|  | **a. Sessions 1–4: Sign-tracking behaviors** |  |  |  |  |  |  |  |  |
| --- | --- | --- | --- | --- | --- | --- | --- | --- | --- |
|  | Lever Contacts |  |  | Probability Lever |  |  | Latency Lever |  |  |
|  | DF | *F* | *p* | DF | *F* | *p* | DF | *F* | *p* |
| *Session* | 3,137.409 | 32.727 | **<.001** | 3,73.415 | 25.881 | **<.001** | 3,59.093 | 29.867 | **<.001** |
| *Treatment* | 1,50.676 | 0.894 | .349 | 1,50.507 | 0.961 | .331 | 1,48.867 | 1.043 | .312 |
| *Phenotype* | 2,50.676 | 27.831 | **<.001** | 2,50.507 | 40.353 | **<.001** | 2,48.867 | 34.499 | **<.001** |
| *Session*×*Treatment* | 3,137.409 | 0.152 | .928 | 3,73.415 | 0.774 | .512 | 3,59.093 | 0.813 | .492 |
| *Session*×*Phenotype* | 6,137.409 | 18.003 | **<.001** | 6,73.415 | 13.343 | **<.001** | 6,59.093 | 16.300 | **<.001** |
| *Treatment* * *Phenotype* | 2,50.676 | 1.040 | .361 | 2,50.507 | 0.923 | .404 | 2,48.867 | 1.127 | .332 |
| *Session*×*Treatment*×*Phenotype* | 6,137.409 | 0.237 | .964 | 6,73.415 | 2.135 | .059 | 6,59.093 | 1.053 | .401 |
|  | **b. Sessions 5–7: Sign-tracking behaviors** |  |  |  |  |  |  |  |  |
|  | Lever Contacts |  |  | Probability Lever |  |  | Latency Lever |  |  |
|  | DF | *F* | *p* | DF | *F* | *p* | DF | *F* | *p* |
| *Session* | 2,70.114 | 0.364 | .696 | 2,95.565 | 1.123 | .329 | 2,72.946 | 0.551 | .579 |
| *Treatment* | 1,48.057 | 0.009 | .923 | 1,49.042 | 0.036 | .850 | 1,48.227 | 0.686 | .412 |
| *Phenotype* | 2,48.057 | 22.509 | **<.001** | 2,49.042 | 26.715 | **<.001** | 2,48.227 | 30.622 | **<.001** |
| *Session*×*Treatment* | 2,70.114 | 0.254 | .776 | 2,95.565 | 0.173 | .841 | 2,72.946 | 0.058 | .944 |
| *Session*×*Phenotype* | 4,70.114 | 1.636 | .175 | 4,95.565 | 1.160 | .333 | 4,72.946 | 1.710 | .157 |
| *Treatment* * *Phenotype* | 2,48.057 | 0.038 | .962 | 2,49.042 | 0.019 | .981 | 2,48.227 | 0.091 | .914 |
| *Session*×*Treatment*×*Phenotype* | 4,70.114 | 0.551 | .699 | 4,95.565 | 0.890 | .473 | 4,72.946 | 0.274 | .894 |
|  | **c. Pretest vs Test: Sign-tracking behaviors in intermediate responders** |  |  |  |  |  |  |  |  |
|  | Lever Contacts |  |  | Probability Lever |  |  | Latency Lever |  |  |
|  | DF | *F* | *p* | DF | *F* | *p* | DF | *F* | *p* |
| *Session* | 1,16 | 3.650 | .074 | 1,16 | 2.767 | .116 | 1,16 | 3.829 | .068 |
| *Treatment* | 1,16 | 0.000 | .984 | 1,16 | 0.038 | .847 | 1,16 | 0.056 | .816 |
| *Session*×*Treatment* | 1,16 | 0.400 | .536 | 1,16 | 0.008 | .928 | 1,16 | 0.321 | .579 |

**Table 3.** Statistical analyses for effects of treatment and phenotype on goal-tracking behaviors. Data from linear mixed effects model analyses for (a) sessions 1–4 and (b) sessions 5–7, with comparisons by treatment and phenotype. A two-way repeated measures ANOVA assessing the effect of treatment within intermediate responders during (c) pretest vs test sessions. Entries, probability, and latency are represented for goal-tracking behaviors. Significant effects and interactions are bolded. These data correspond to that shown in Figure 4.

| **a. Sessions 1–4: Goal-tracking behaviors** |  |  |  |  |  |  |  |  |  |
| --- | --- | --- | --- | --- | --- | --- | --- | --- | --- |
|  | Food Cup Entries |  |  | Probability Food Cup |  |  | Latency Food Cup |  |  |
|  | DF | *F* | *p* | DF | *F* | *p* | DF | *F* | *p* |
| *Session* | 3,76.837 | 7.035 | **<.001** | 3,95.188 | 18.529 | **<.001** | 3,86.212 | 17.968 | **<.001** |
| *Treatment* | 1,51.163 | 0.278 | .601 | 1,50.286 | 0.022 | .883 | 1,51.460 | 0.001 | .972 |
| *Phenotype* | 2,51.163 | 4.666 | **.014** | 2,50.286 | 5.562 | **.007** | 2,51.460 | 7.078 | **.002** |
| *Session*×*Treatment* | 3,76.837 | 0.284 | .837 | 3,95.188 | 0.260 | .854 | 3,86.212 | 0.068 | .977 |
| *Session*×*Phenotype* | 6,76.837 | 11.583 | **<.001** | 6,95.188 | 19.421 | **<.001** | 6,86.212 | 17.423 | **<.001** |
| *Treatment*×*Phenotype* | 2,51.163 | 0.409 | .666 | 2,50.286 | 0.056 | .946 | 2,51.460 | 0.303 | .740 |
| *Session*×*Treatment*×*Phenotype* | 6,76.837 | 0.665 | .678 | 6,95.188 | 1.562 | .167 | 6,86.212 | 0.881 | .512 |
| **b. Sessions 5–7: Goal-tracking behaviors** |  |  |  |  |  |  |  |  |  |
|  | Food Cup Entries |  |  | Probability Food Cup |  |  | Latency Food Cup |  |  |
|  | DF | *F* | *p* | DF | *F* | *p* | DF | *F* | *p* |
| *Session* | 2,64.226 | 0.203 | .817 | 2,95.255 | 0.812 | .447 | 2,69.999 | 1.615 | .206 |
| *Treatment* | 1,48.429 | 4.349 | **.042** | 1,48.154 | 7.561 | **.008** | 2,48.804 | 3.945 | .053 |
| *Phenotype* | 2,48.436 | 32.846 | **<.001** | 2,48.161 | 57.539 | **<.001** | 2,48.811 | 56.982 | **<.001** |
| *Session*×*Treatment* | 2,64.226 | 0.563 | .572 | 2,95.255 | 0.579 | .563 | 2,69.999 | 0.083 | .920 |
| *Session*×*Phenotype* | 4,64.283 | 0.451 | .771 | 4,95.255 | 0.097 | .983 | 4,70.005 | 0.234 | .918 |
| *Treatment*×*Phenotype* | 2,48.436 | 1.426 | .250 | 2,48.161 | 4.886 | **.012** | 2,48.811 | 2.655 | .080 |
| *Session*×*Treatment*×*Phenotype* | 4,64.283 | 1.742 | .152 | 4,95.255 | 1.752 | .145 | 4,70.005 | 1.938 | .114 |
| **c. Pretest vs Test: Goal-tracking behaviors in intermediate responders** |  |  |  |  |  |  |  |  |  |
|  | Food Cup Entries |  |  | Probability Food Cup |  |  | Latency Food Cup |  |  |
|  | DF | *F* | *p* | DF | *F* | *p* | DF | *F* | *p* |
| *Session* | 1,16 | 0.035 | .853 | 1,16 | 0.016 | .901 | 1,16 | 0.165 | .690 |
| *Treatment* | 1,16 | 5.560 | **.031** | 1,16 | 5.149 | **.037** | 1,16 | 2.998 | .103 |
| *Session*×*Treatment* | 1,16 | 4.448 | .051 | 1,16 | 8.311 | **.011** | 1,16 | 4.105 | .060 |

### Effects of inhibiting the LH–PVT pathway on PavCA behavior (Test)

#### Inhibition of the LH–PVT pathway has no impact on sign-tracking

Chemogenetic inhibition of the LH–PVT pathway via systemic CNO administration had no effect on the expression of sign-tracking behaviors (Table 1b, 2b). There were no significant differences in the number of lever-cue contacts (Figure 3a, Figure 4a), probability to approach the lever-cue (Figure 3c, Figure 4c), or latency to approach the lever-cue (Figure 3e, Figure 4e) between VEH- and CNO-treated rats on sessions 5–7. This was true when comparing treatment groups collapsed across phenotypes (Table 1b; Figure 3a, c, e) and when phenotype was included as a variable (Table 2b; Figure 4a, c, e). As expected, sign-tracking behavior changed across sessions, and there was a significant effect of phenotype for each lever-directed measure (Table 2b). *Post hoc* comparisons revealed that STs had significantly more lever-cue contacts (*p* < 0.001; Figure 4a), a higher probability to approach the lever-cue (*p* < 0.001; Figure 4c), and a faster latency to approach the lever-cue (*p* < 0.001; Figure 4e) relative to IRs and GTs on sessions 5–7. Thus, LH–PVT pathway inhibition via DREADD activation had no effect on sign-tracking behavior.

To further assess differences in behavior as a function of LH–PVT pathway inhibition, we directly compared sign-tracking behaviors on session 4 (pretest) to the average of sessions 5–7 (test) (Table 1c, 2c). There was a significant effect of session for all lever-directed behaviors (Table 1c; Figure 3b, d, f). *Post hoc* comparisons revealed that rats made more lever-cue contacts (*p* = 0.008; Figure 3b), had a higher probability to approach the lever-cue (*p* = 0.005; Figure 3d), and had a faster latency to approach the lever-cue (*p* = 0.002; Figure 3f) during test sessions relative to pretest. This continued change in responding is expected with subsequent training sessions (Meyer et al., 2012). Consistent with these findings, sign-tracking behavior was not altered due to LH–PVT inhibition when directly comparing pretest to test sessions within phenotypes (Table 2c; Figure 4b, d, f; *data not shown for ST or GT*).

#### Inhibition of the LH–PVT pathway decreases goal-tracking behavior

Chemogenetic inhibition of the LH–PVT pathway decreased goal-tracking behaviors, and this reduction was phenotype-dependent. When collapsed across phenotype, there was a significant effect of treatment for goal-tracking behavior on sessions 5–7 (Table 1e). Relative to VEH-treated rats, CNO-treated rats had decreased food cup entries when the lever-cue was presented (effect of treatment: *F*_1,52.137_ = 5.147, *p* = 0.027; Figure 3g), decreased probability to approach the food cup (effect of treatment: *F*_1,52.025_ = 5.119, *p* = 0.028; Figure 3i), and a trend for an increased latency to approach the food cup (effect of treatment: *F*_1,52.012_ = 3.655, *p* = 0.061; Figure 3k). Thus, inhibition of the LH–PVT pathway decreased the expression of goal-tracking behavior. Together with the lack of effect on sign-tracking, these data suggest that LH–PVT inhibition selectively reduced food cup-directed conditioned responding.

When phenotype was considered as a variable, there was a significant effect of phenotype (*p* < 0.001) and treatment (*p* ≤ 0.05) for all measures of goal-tracking behavior over sessions 5–7 (Table 3b), and a significant phenotype × treatment interaction for the probability to approach the food cup (*F*_2,48.436_ = 4.886, *p* = 0.012). This interaction indicated that the treatment effect was not evenly distributed across phenotypes. As expected, GTs showed more robust responding directed toward the food cup relative to STs, and this was independent of treatment (Table 3b; Figure 4g, i, k). As shown in Figure 4i, however, VEH-treated IR rats showed a pattern of food cup approach more similar to GTs, whereas CNO-treated IR rats showed reduced food cup approach and a pattern more similar to STs by the end of the testing phase. In agreement, *post hoc* comparisons revealed that CNO-treated IRs had a significantly reduced probability to approach the food cup relative to VEH-treated IRs (*p* < 0.001, Cohen’s *d* = 1.53). Further, within treatment groups, VEH-treated IRs had a significantly higher probability to approach the food cup relative to VEH-treated STs (*p* < 0.001, Cohen’s *d* = 2.48), but not VEH-treated GTs (*p* = 0.372); whereas CNO-treated IRs had a significantly lower probability to approach the food cup relative to CNO-treated GTs (*p* < 0.001, Cohen’s *d* = 2.65), but not CNO-treated STs (*p* = 0.063). Taken together, inhibition of the LH–PVT pathway significantly and selectively decreased the probability to goal-track in IR rats.

We also compared session 4 (pretest) to the average of sessions 5–7 (test) for goal-tracking behaviors in VEH- and CNO-treated rats (Table 1f, Figure 3h, j, l). As shown in Figure 3j, there was a significant session × treatment interaction for probability to approach the food cup when the lever-cue was presented (*F*_1,52_ = 10.532, *p* = 0.002). *Post hoc* comparisons revealed that CNO-treated rats had a significantly higher probability to approach the food cup on the pretest session relative to the test session (*p* = 0.005, Cohen’s *d* = 0.35). Additionally, on the test session, CNO-treated rats had a significantly lower probability to approach the food cup relative to VEH-treated rats (*p* = 0.028, Cohen’s *d* = 0.15). Thus, the pretest–test comparison supported the session 5–7 analysis, showing that LH–PVT inhibition reduced the probability of food cup approach during cue presentation.

We further compared goal-tracking behaviors between pretest and test sessions in IRs (Table 3c). As shown in Figure 4j, there was a significant session × treatment interaction (*F*_1,16_ = 8.311, *p* = 0.011) for probability to approach the food cup. *Post hoc* comparisons revealed that CNO-treated IRs had a significantly higher probability to approach the food cup on the pretest session relative to the test session (*p* = 0.042, Cohen’s *d* = 0.68). Additionally, on the test session, CNO-treated IR rats had a significantly lower probability to approach the food cup relative to VEH-treated IR rats (*p* = 0.006, Cohen’s *d* = 0.36). Taken together, these data support those above, demonstrating that LH–PVT inhibition attenuates the probability to goal-track, and does so selectively in IR rats.

#### Inhibition of the LH–PVT pathway does not alter food cup entries during the inter-trial interval

To determine whether LH–PVT inhibition reduced food cup-directed behavior more broadly, we analyzed food cup entries during the inter-trial interval (ITI), when the lever-cue was not present. There was an overall decrease in ITI food cup entries across sessions 1–4 (effect of session, F_3,92.361_ = 5.794, *p* = 0.001) and 5–7 (effect of session, F_2,97.023_ = 15.236, *p* < 0.001) (Extended Data Figure 2a; Extended Data Table 2a, b), as expected with learning of the cue-reward relationship. There was no effect of treatment on ITI food cup entries during sessions 5–7 when rats were analyzed with phenotypes collapsed (Extended Data Figure 2a; Extended Data Table 2b); and this was also true when phenotype was included as a variable (Extended Data Figure 2b–d; Extended Data Table 2b). Although there was a significant session × treatment × phenotype interaction for food cup entries during sessions 5–7 (F_4,48.007_ = 3.251, *p* = 0.019), follow-up comparisons showed no significant VEH–CNO differences within STs, IRs, or GTs (Extended Data Table 2c). Since food cup entries during the ITI were not affected by treatment either in the total population or in IRs, LH–PVT inhibition appears to selectively reduce cue-evoked goal-tracking behavior without a general reduction in food cup entries. In agreement, all rats consumed all of the food pellets that were delivered during PavCA sessions.

#### Inhibition of the LH–PVT pathway during PavCA does not affect the conditioned reinforcing properties of the lever-cue

Prior treatment with CNO during PavCA did not affect performance on the subsequent conditioned reinforcement test, which was conducted in the absence of acute VEH or CNO treatment. Thus, LH–PVT inhibition during PavCA did not alter the conditioned reinforcing properties of the lever-cue under these conditions (Extended Data Figure 3, Extended Data Table 3).

#### CNO in the absence of DREADD does not impact behavior

The purpose of Experiment 2 was to confirm that the effects reported above were DREADD-mediated; therefore, we assessed the effects of 5 mg/kg CNO (i.p.) in the absence of DREADD receptors in the brain. CNO-treated rats were no different from VEH-treated rats in sign-tracking or goal-tracking behaviors during PavCA sessions 1–4 or sessions 5–7 (Extended Data Figure 4; Extended Data Table 4), and the same was true when phenotype was considered as a variable (Extended Data Figure 4; Extended Data Tables 5, 6). Although there was a significant session × treatment × phenotype interaction for food cup entries during sessions 5–7 (F_4,110.272_ = 3.112, *p* = 0.018; Extended Data Table 6b), follow-up analyses within phenotype did not reveal significant session × treatment interactions in STs, IRs, or GTs (Extended Data Table 6c). Thus, CNO did not produce a treatment-dependent change in food cup entries in rats without DREADD expression. Further, prior VEH or CNO treatment did not alter the conditioned reinforcing properties of the lever-cue (Extended Data Figure 5, Extended Data Table 7). Thus, while STs, IRs, and GTs showed the expected behavioral patterns, VEH- and CNO-treated rats did not differ within any phenotype in the absence of DREADD. The CNO-elicited effects reported above, therefore, were dependent on DREADD expression.

## Discussion

We examined the role of the LH–PVT pathway in the expression of conditioned responses in a Pavlovian conditioned approach paradigm. Chemogenetic inhibition of this pathway decreased goal-tracking behavior, with effects confined to intermediate responders (IRs) when the PavCA phenotype was considered. Thus, LH–PVT signaling supports goal-tracking behavior in a phenotype-dependent manner. Notably, all rewards were consumed during PavCA sessions, and LH–PVT inhibition did not impact food cup entries during the inter-trial interval. Thus, the reduction in goal-tracking appears specific to cue-evoked food cup-directed responding, rather than reflecting disrupted reward consumption or a general reduction in food cup-directed behavior.

The selective impact of LH–PVT inhibition on IRs is consistent with the idea that once a prepotent conditioned response is established, it can be difficult to disrupt, particularly in animals at the extremes of the population (Campus et al., 2019; Flagel et al., 2009). IRs display flexibility in their responding, vacillating between cue- and food cup-directed behaviors, and may therefore be especially sensitive to manipulations that bias response selection. This interpretation aligns with evidence that the PVT integrates cortical and hypothalamic signals to shape cue-reward processing and reward seeking (Otis et al., 2019), and may arbitrate motivational competition when multiple responses are available (Choi et al., 2019; McNally, 2021). The IR-specific effect was most apparent in the probability of food cup approach, such that LH–PVT inhibition reduced the likelihood that IR rats initiated a goal-tracking response during cue presentation. Measures of response vigor and approach latency were less affected, suggesting an effect on response selection rather than general performance. Because most prior work has emphasized sign-tracker and goal-tracker extremes (Campus et al., 2019; Flagel et al., 2007; Haight et al., 2020; Robinson & Flagel, 2009), the circuit mechanisms supporting IR behavior remain poorly characterized. The current findings identify the LH–PVT pathway as a potential leverage point for the flexible cue-motivated responding seen in IRs, a behavioral pattern that may better capture population-level variability.

The absence of an effect on sign-tracking may appear to diverge from work emphasizing bottom-up contributions to incentive-salience attribution in sign-trackers. Prior studies implicate hypothalamic–thalamic–striatal circuitry in incentive motivational processes (Flagel, Cameron, et al., 2011; Haight et al., 2017; Haight et al., 2020), making it reasonable to predict that suppressing LH–PVT signaling would attenuate sign-tracking. However, our data indicate that broad LH–PVT inhibition is not sufficient to disrupt the expression of sign-tracking behavior. One possibility is that sign-tracking depends on specific components of the LH–PVT projection, particularly orexinergic transmission, rather than global pathway activity. Consistent with this, orexin receptor antagonism in the PVT reduces the incentive value of a food-paired cue in sign-trackers (Haight et al., 2020). Additionally, LH–PVT signaling may be more critical during acquisition than expression of conditioned responses. Supporting this possibility, excitotoxic lesions of the LH performed before learning reduced subsequent sign-tracking behavior (Haight et al., 2020). Moreover, posterior PVT activity gates associative learning during acquisition but not expression (Zhu et al., 2018). In head-fixed mice, optogenetic silencing of the PVT during cue (CS) presentation or reward (US) consumption weakened CS–US learning, reflected by reduced anticipatory licking, but only when delivered during acquisition (Zhu et al., 2018). Together, these findings motivate targeted tests of cell-type- and phase-specific mechanisms (acquisition vs. expression) within LH–PVT circuitry.

A key limitation of the present approach is that the viral manipulation was not cell-type specific and likely engaged multiple LH–PVT systems. LH–PVT projections are heterogeneous, including dense orexinergic, GABAergic, and glutamatergic inputs (Jennings et al., 2015; Kirouac et al., 2005; Li & Kirouac, 2012; Otis et al., 2019), and the PVT itself contains molecularly distinct neuronal subtypes (Gao et al., 2023). Accordingly, distinct LH–PVT populations likely contribute to reward-related behavior via different mechanisms. Orexinergic signaling in the PVT promotes appetitive motivation (Barson et al., 2015; Li et al., 2009; Stratford & Wirtshafter, 2013), including sign-tracking (Haight et al., 2020). In parallel, LH–PVT GABAergic projections are substantial (Otis et al., 2019), and their stimulation elicits consummatory and reward-seeking behaviors (Wu et al., 2015; Zhang & van den Pol, 2017); whereas optogenetic inhibition of LH GABA neurons after learning reduces motivation to seek reward without altering consumption (Sharpe et al., 2017). Moreover, distinct LH GABAergic subpopulations appear to differentially track reward seeking versus consumption (Jennings et al., 2015). This is relevant to our findings, which show reduced goal-tracking without changes in reward consumption, suggesting that LH–PVT inhibition disrupted cue-evoked approach rather than primary reinforcement or consummatory processes. However, because feeding was not assessed outside of PavCA, the absence of a consumption effect here should not be taken as evidence that LH–PVT signaling is uninvolved in feeding more broadly (Hoebel & Teitelbaum, 1962; Margules & Olds, 1962). Consistent with broader feeding roles, the LH is functionally heterogeneous and can support both food intake and self-stimulation depending on stimulation site and experience (Urstadt & Berridge, 2020). Further, the PVT is embedded in feeding-related circuitry via connections to the NAc shell (Stratford, 2005; Stratford, 2007) and hypothalamus (Chen & Su, 1990; Moga et al., 1995; Stratford, 2005; Stratford, 2007), and activation of GABA_A_ receptors in the PVT dose-dependently increases food intake in sated rats when food is freely available (Stratford & Wirtshafter, 2013). Methodological differences (e.g., systemic CNO inhibition of LH–PVT neurons during PavCA versus local manipulation of PVT GABA_A_ signaling on a feeding test) likely also contribute to divergent outcomes across studies. This point is further supported by evidence that orexin receptor 1 (OX1R)–expressing PVT neurons are activated in response to food-paired cues (Choi et al., 2010), and GABAergic LH–PVT inputs onto glutamatergic PVT–NAc neurons have been shown to mediate consummatory behavior, such as licking (Otis et al., 2019). These distinctions may help explain why LH–PVT manipulations affect reward seeking and consummatory behavior differently across paradigms and under conditions that vary satiety state and food availability.

Anatomical specificity is also important to consider, as mCherry-positive cell body expression was consistently observed in the ZI, which is adjacent to the LH and difficult to fully avoid with the current viral targeting approach. ZI–PVT projections have been implicated in food seeking and consumption, including motivational vigor during food seeking and rapid binge-like eating following stimulation of ZI GABA neurons or their projections to the PVT (Ye et al., 2023; Zhang & van den Pol, 2017). Although we cannot fully dissociate contributions of LH–PVT neurons from adjacent ZI–PVT neurons in the current study, the data described above argue against a broad disruption of feeding or general appetitive behavior. Instead, the present findings suggest that inhibition of this lateral hypothalamic/ZI region projecting to the PVT selectively altered conditioned food cup-directed responding during cue presentation. Dissociating LH and ZI contributions to PVT-dependent cue-motivated behavior will require more spatially restricted and cell-type-specific approaches.

To date, PVT-focused studies within the sign-tracker/goal-tracker model have largely targeted the entire PVT rather than isolating anterior or posterior subregions (Campus et al., 2019; Haight et al., 2015; Haight et al., 2017; Kuhn et al., 2022; Kuhn et al., 2018). However, the anterior PVT (aPVT) and posterior PVT (pPVT) have distinct functions. Generally, the aPVT appears to regulate reward-seeking and arousal (Cheng et al., 2018; Do-Monte et al., 2017; Gao et al., 2020; Kolaj et al., 2012), whereas the pPVT is largely involved in stress and anxiety (Barson & Leibowitz, 2015; Bhatnagar & Dallman, 1999; Li et al., 2010a, 2010b). Because we targeted the entire rostro-caudal extent of the PVT, it is difficult to determine whether the observed effects arise primarily from aPVT, pPVT, or their interaction. Further supporting this point, recent work shows that parallel PVT projections to the NAc can dissociably encode features of goal-oriented behavior, including the execution and termination of goal-directed actions (Beas et al., 2024). This is especially relevant given that LH inputs, including orexinergic projections, may differentially engage anterior and posterior PVT subregions, which could have distinct consequences for cue-motivated behavior. Selective targeting of the aPVT and pPVT would clarify whether LH–PVT projections regulate distinct features of PavCA behavior through subregion-specific PVT circuits. Because orexinergic LH–PVT projections have been implicated in incentive salience attribution (Haight et al., 2020; Haight et al., 2017), orexinergic LH–aPVT projections may be a particularly important target for future studies in this model.

Together with our findings, recent work supports viewing the PVT and LH–PVT pathway as a collection of specialized, state-dependent circuits rather than a single functional unit. This view is consistent with the early framework proposed by Kelley and colleagues, in which the PVT was positioned as a critical node in a hypothalamic-thalamic-striatal circuit that integrates state-related signals to guide motivated behaviors (Kelley et al., 2005). More recent work suggests that PVT contributions to motivated behavior are best understood through projection-defined and state-dependent circuit mechanisms, rather than through whole-region manipulations alone (Beas et al., 2024). Consistent with this view, distinct PVT–NAc circuits may differentially encode valence and salience (Rivera-Irizarry et al., 2023), while PVT dopamine D2R+ neurons integrate hunger, thirst, and cue-related information to guide motivated behavior (Machen et al., 2026). Collectively, these findings suggest that the behavioral effects observed here may depend on the specific LH inputs engaged, the PVT subregion and output pathway recruited, and the motivational state of the animal. Whether LH–PVT inhibition regulates PavCA behavior similarly across internal states such as satiety, food restriction, stress, or altered arousal remains an open question.

Although inclusion of both sexes was intended for this work, we were unable to add females to the study due to the COVID-19 pandemic and subsequent resource constraints. Thus, the current findings are limited to male rats. It remains unclear whether LH–PVT inhibition would produce similar effects in females. This limitation warrants consideration, as sex differences have been reported in PavCA behavior (Hughson et al., 2019; Stringfield et al., 2019) and in neural systems that regulate reward seeking and motivated behavior (Becker & Koob, 2016; Becker et al., 2017). However, the direction of these effects is difficult to predict, as sex differences in PavCA behavior appear to depend on experimental conditions, vendor/strain, behavioral measure, and exposure history (Bien & Smith, 2023; Turfe et al., 2024). Future work should include both males and females to determine whether the LH–PVT pathway regulates goal-tracking behavior and behavioral flexibility similarly across sex.

Sample sizes in the current study were limited by uneven phenotype distributions across vendor/barrier cohorts and by stringent anatomical inclusion criteria for DREADD expression. This resulted in uneven group sizes, particularly within phenotype-specific comparisons. Although this limitation may reduce sensitivity to detect smaller effects, the primary IR effect was large, particularly for probability to approach the food cup (Cohen’s *d* = 1.53). In contrast, variability within the ST and GT treatment groups was low, suggesting that the absence of treatment effects in these phenotypes was not simply due to increased variance in behavioral responding.

In the current study, we targeted afferents from the LH to the PVT to elucidate the role of this pathway in phenotypically distinct cue-motivated behaviors. CNO administration in non-DREADD rats had no off-target effects in PavCA or CRT, supporting the interpretation that the behavioral effects observed here were DREADD-mediated. Together, these data indicate that LH–PVT pathway inhibition reduces reward-directed conditioned responding in behaviorally flexible PavCA phenotypes without broadly disrupting sign-tracking, ITI food cup entries, or reward consumption within the texting context. Future studies should combine cell-type-specific manipulations within the LH with subregion-specific targeting of the PVT to dissociate contributions of orexinergic and non-orexinergic signaling. Collectively, these findings refine the role of the LH–PVT pathway in cue-motivated behavior and suggest that this circuit is especially relevant when reward-directed conditioned responses are flexible rather than strongly biased toward the cue or reward location.

## Author Contributions

A.G.I, P.C., and S.B.F. designed research. A.G.I., J.K.B., A.E.T., S.C., and J.L. performed research. A.G.I. and S.B.F. analyzed data. S.C. edited the paper, A.G.I. and S.B.F. wrote the paper.

## Acknowledgements

Thank you to members of the Flagel Lab for their support and feedback on these studies.

Authors report no conflict of interest.

## Extended Data

**Extended Data Figure 1.**
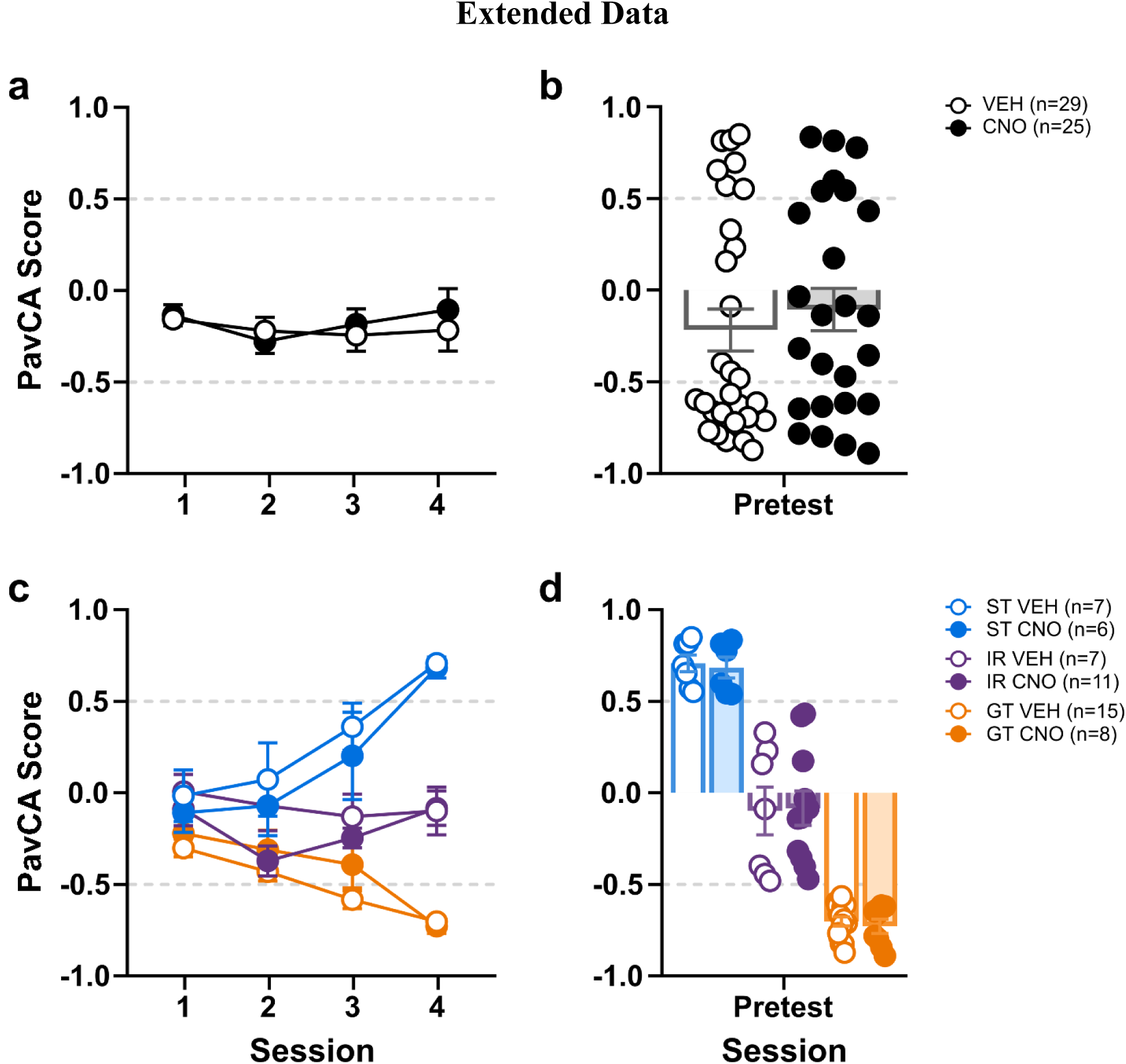
Acquisition of Pavlovian Conditioned Approach (PavCA) prior to treatment. Data are expressed as mean ± SEM for PavCA Score (a,c) over sessions 1–4 and (b,d) on session 4 (pretest). There are no significant differences between treatment or phenotype/treatment groups in (a,c) PavCA Score across sessions 1–4 or (b,d) PavCA Index on the pretest session. (c,d) There are the expected differences between phenotypes for PavCA Score such that STs have a higher score than IRs and GTs by session 4.

**Extended Data Figure 2.**
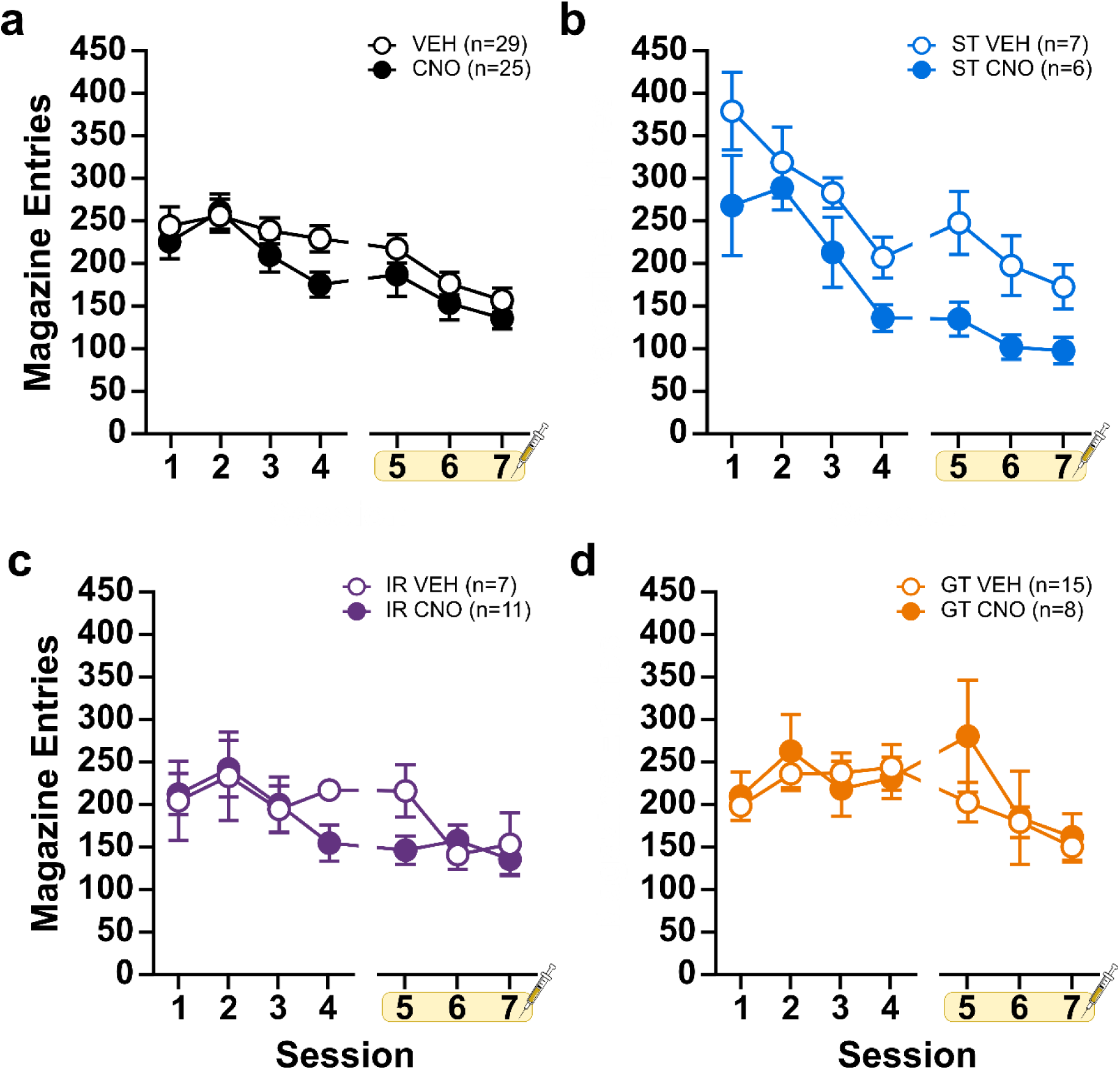
Chemogenetic inhibition of the LH–PVT pathway does not impact food cup entries during the inter-trial interval. Data are expressed as mean ± SEM for food cup entries during the inter-trial interval (ITI) within Pavlovian conditioned approach (PavCA). Food cup entries during the ITI period across sessions 1–7 (a) between treatment groups collapsed and (b–d) with treatment and phenotype groups expanded. On sessions 5–7, rats received either VEH or CNO prior to testing. There were no treatment effects on food cup entries during the ITI period (a) between treatment groups or (b–d) within any of the phenotypes for sessions 5–7.

**Extended Data Figure 3.**
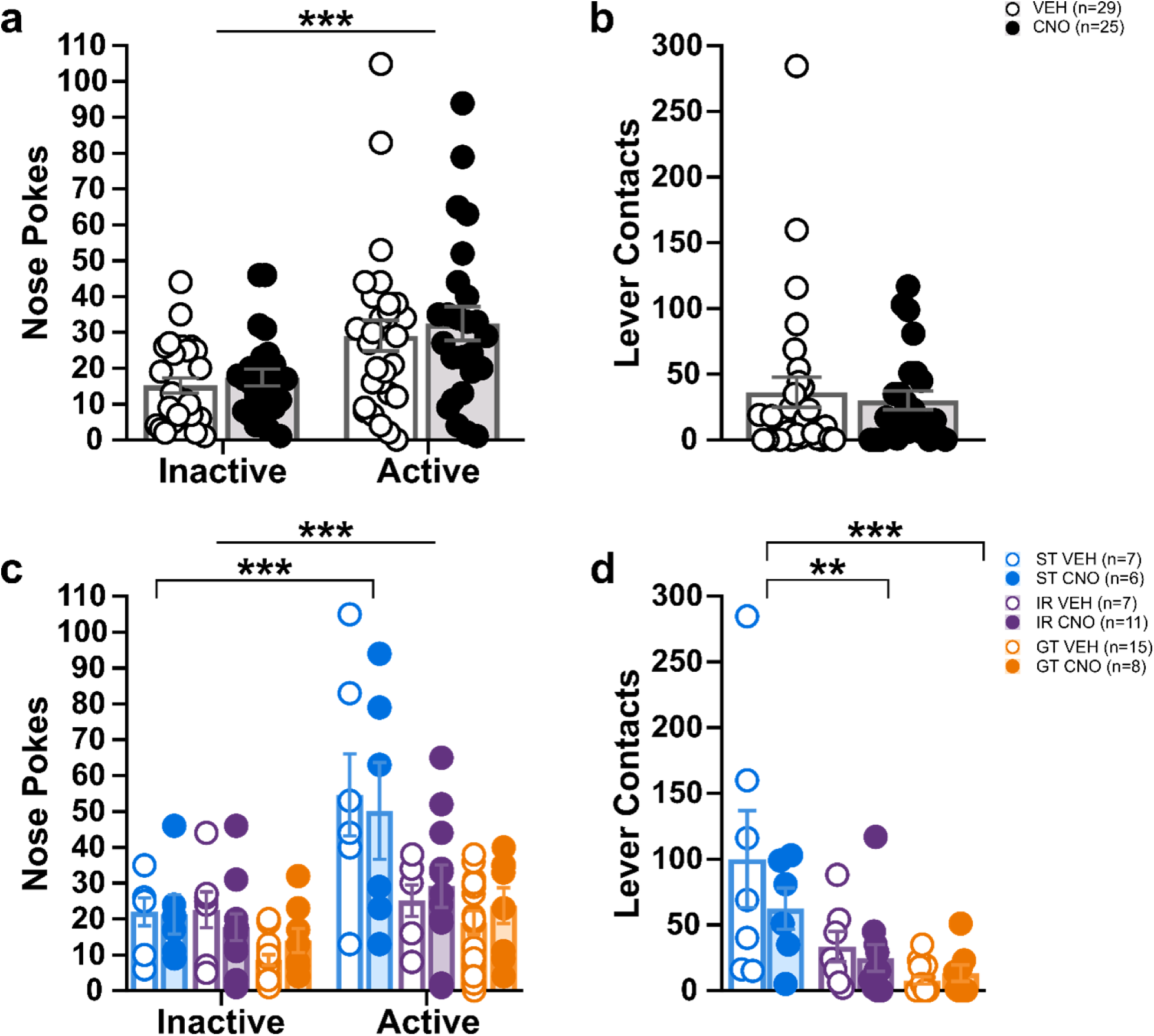
Prior chemogenetic inhibition of the LH–PVT pathway does not impact the conditioned reinforcing properties of the lever-cue. Data are expressed as mean ± SEM for (a,c) nose pokes in the inactive and active ports and (b,d) lever contacts during the conditioned reinforcement test (CRT). There are no effects of treatment in the CRT. (a,c) Rats nose poke in the active port more than the inactive port. (c) STs have more active than inactive nose pokes; and they have more active nose pokes than IRs and GTs. (d) STs have more lever contacts than IRs and GTs. Straight line indicates a significant effect of port. Brackets indicate significant differences between phenotypes or ports, **p < 0.01 and ***p < 0.001.

**Extended Data Figure 4.**
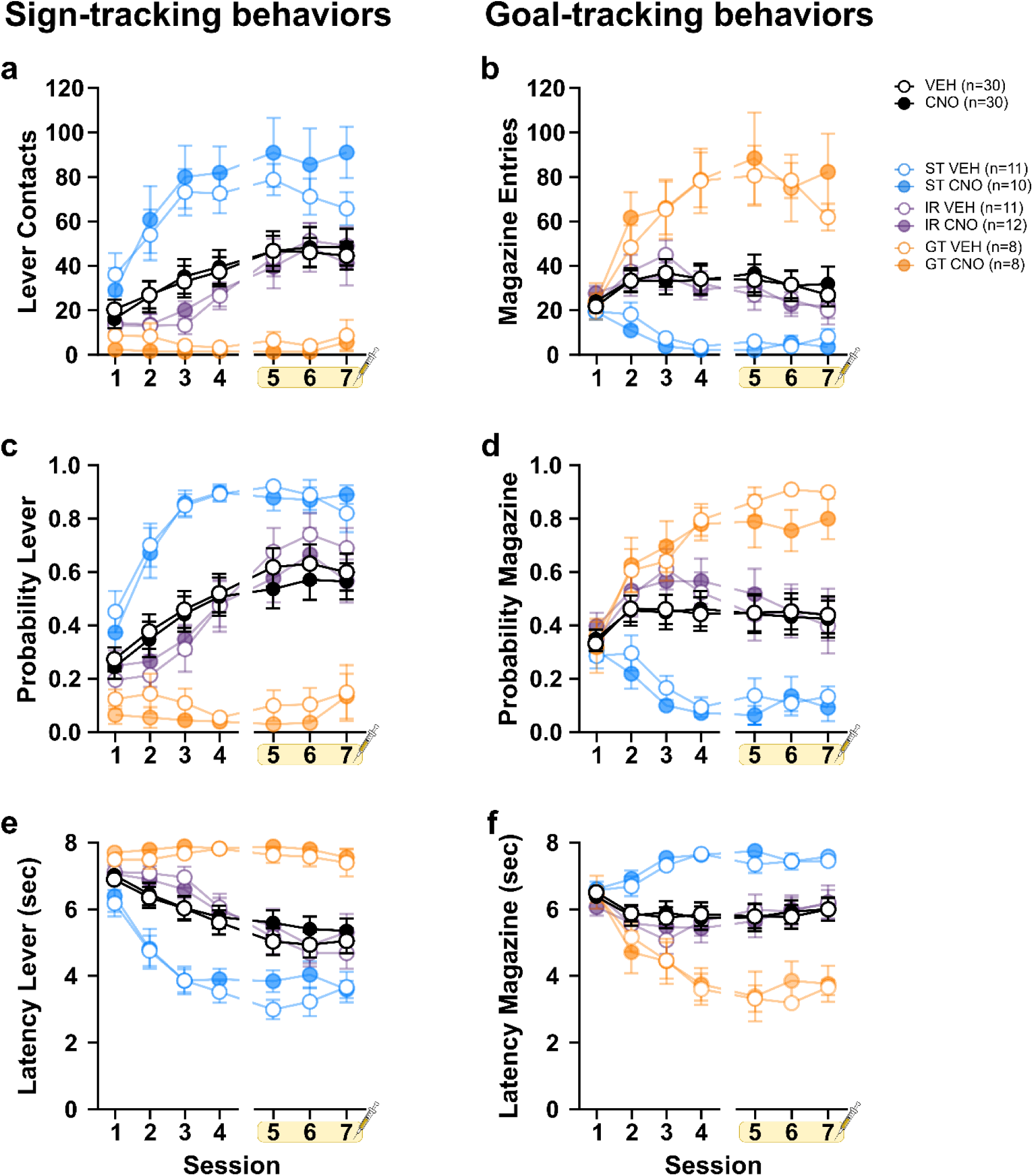
Clozapine-N-oxide has no effects on Pavlovian Conditioned Approach (PavCA) behavior the absence of DREADD receptors. Data are expressed as mean ± SEM for rats that did not receive surgery for DREADD expression during Pavlovian conditioned approach (PavCA). (a–c) Sign-tracking and (d–f) goal-tracking behaviors across sessions 1–7 shown with treatment groups (black lines) collapsed in the foreground and with phenotype/treatment groups expanded in background (muted color). (a–c) Lever- and (d–f) food cup-directed behaviors for (a,d) number of contacts/entries, (b,e) probability to approach, or (c,f) latency to approach. Treatment had no effect on sign- or goal-tracking behaviors.

**Extended Data Figure 5.**
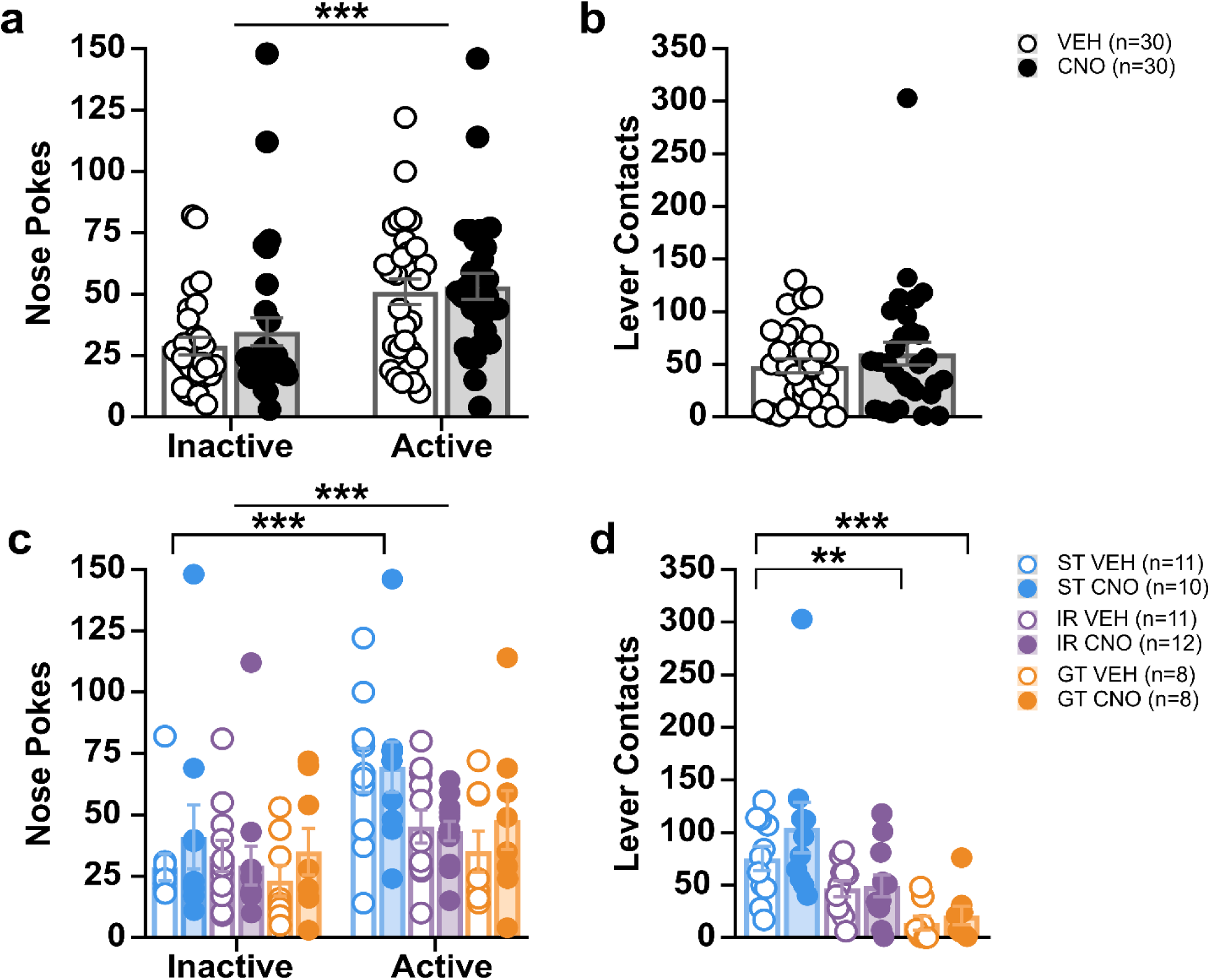
Clozapine-N-oxide has no effects on the conditioned reinforcing properties of the lever in the absence of DREADD expression. Data are expressed as mean ± SEM for rats that did not receive surgery for DREADD expression during (a–d) a conditioned reinforcement test (CRT). Behavior during the CRT is shown as (a–b) treatment groups collapsed and (c–d) treatment and phenotype groups expanded. (a,c) Nose pokes in the inactive and active ports and (b,d) lever contacts. There were no significant effects of treatment on behavior during the CRT. (a,c) Rats nose poke in the active port more than the inactive port. (c) STs have more active than inactive nose pokes. (d) STs have more lever contacts than IRs and GTs. Straight line indicates a significant effect of port. Brackets indicate significant differences between phenotypes or ports, **p < 0.01 and ***p < 0.001.

**Extended Data Table 1.**
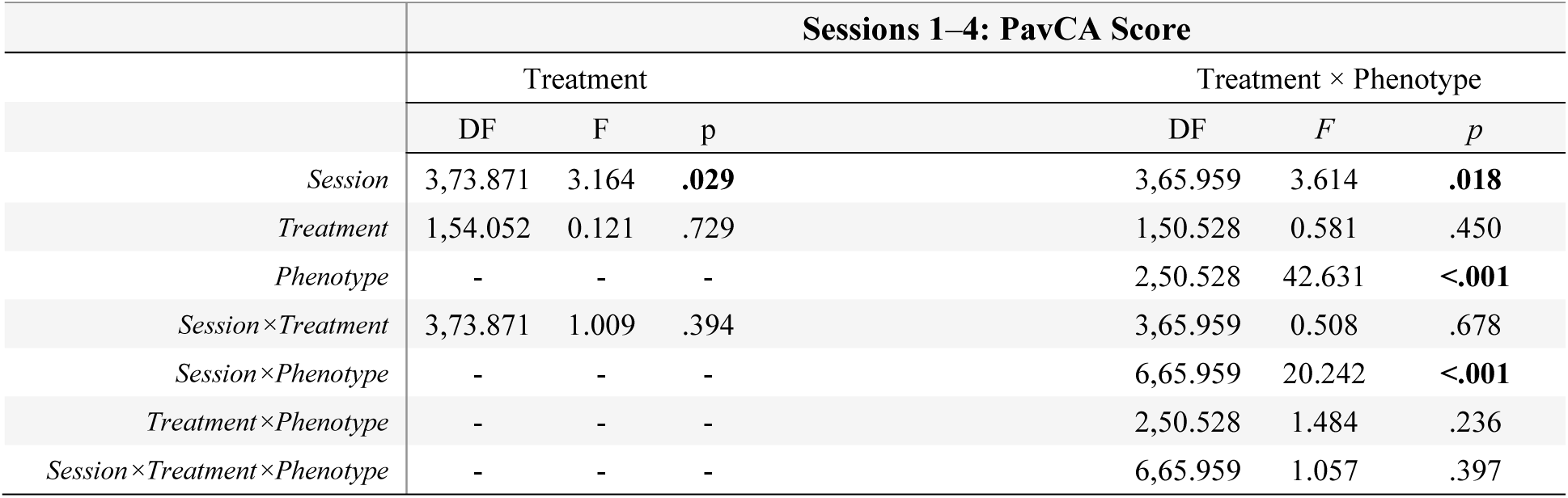
Statistical analyses for PavCA Score across sessions 1–4. Data from linear mixed effects model analyses for PavCA Score on sessions 1–4. The left panel compares rats by treatment, collapsed across phenotype. The right panel compares rats with treatment and phenotype as variables. Significant effects and interactions are bolded. These data correspond to data shown in Extended Data Figure 1.

**Extended Data Table 2.**
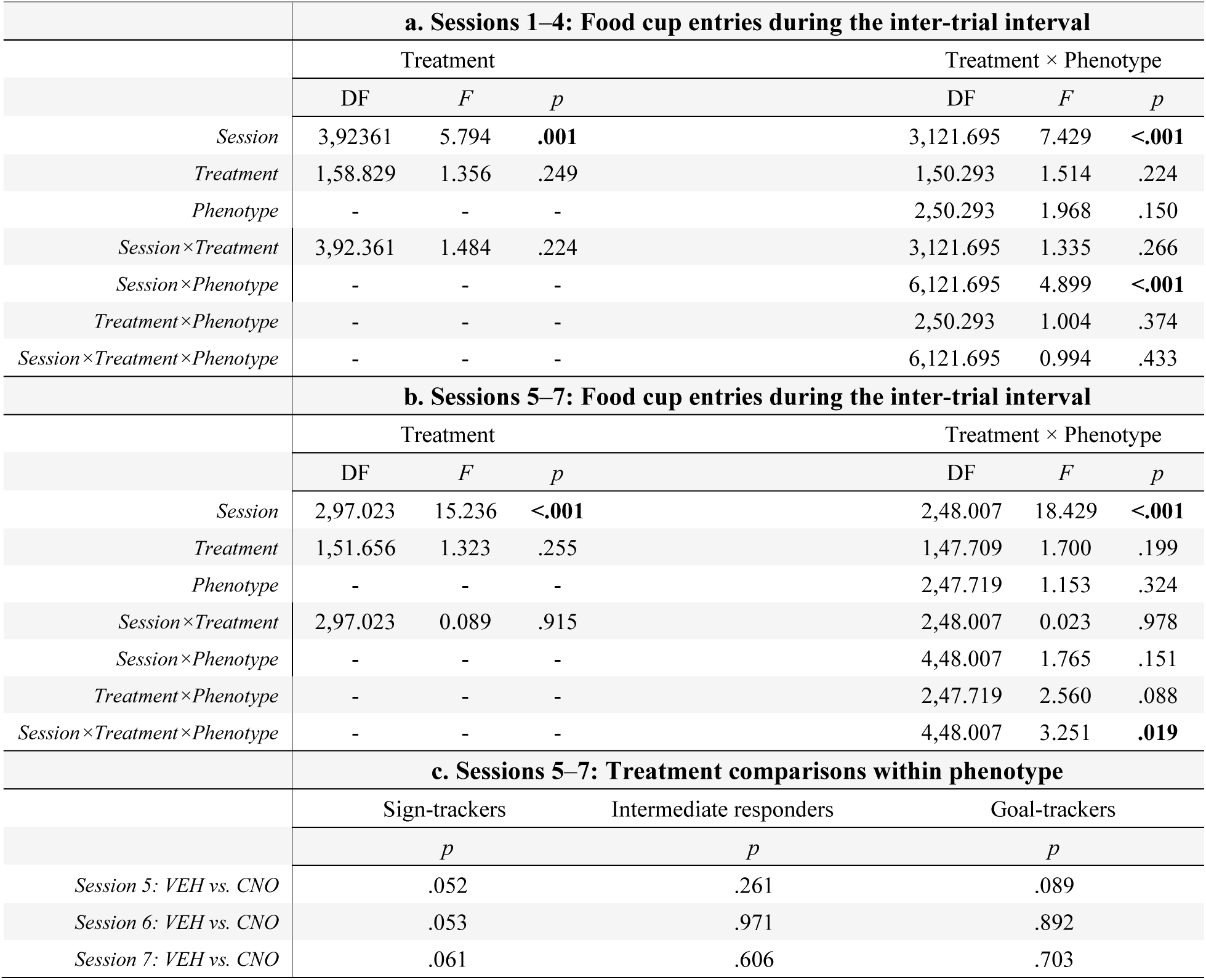
Statistical analyses for the effects of treatment and phenotype on food cup entries during the inter-trial interval. Data are from linear mixed effects model analyses for (a) sessions 1–4 and (b) sessions 5–7; (c) shows follow-up treatment comparisons within phenotype for the significant session × treatment × phenotype interaction. For (a,b), the left panel compares rats by treatment, collapsed across phenotype, and the right panel compares rats with treatment and phenotype as variables. Significant effects and interactions are bolded. These data correspond to data shown in Extended Data Figure 2.

**Extended Data Table 3.** Statistical analyses for behavior during the Conditioned Reinforcement Test (CRT). Data from a mixed ANOVA comparing (left column) nose pokes into each port and a two-way ANOVA comparing lever contacts during the conditioned reinforcement test. Analysis was conducted (a) between treatment groups and (b) with treatment and phenotype as variables. Significant effects and interactions are bolded. These data correspond to the data shown in Extended Data Figure 3.

|  | **a. Session 8: Conditioned Reinforcement** |  |  |  |  |  |
| --- | --- | --- | --- | --- | --- | --- |
|  | Nose Pokes |  |  | Lever Contacts |  |  |
|  | DF | *F* | *p* | DF | *F* | *p* |
| *Port* | 1,52 | 31.889 | **<.001** | - | - | - |
| *Treatment* | 1,52 | 0.433 | .514 | 1,52 | 0.196 | .660 |
| *Port*×*Treatment* | 1,52 | 0.053 | .819 | - | - | - |
|  | **b. Session 8: Conditioned Reinforcement** |  |  |  |  |  |
|  | Nose Pokes |  |  | Lever Contacts |  |  |
|  | DF | *F* | *p* | DF | *F* | *p* |
| *Port* | 1,48 | 44.721 | **<.001** | - | - | - |
| *Treatment* | 1,48 | 0.035 | .853 | 1,48 | 1.282 | .263 |
| *Phenotype* | 2,48 | 9.221 | **<.001** | 2,48 | 11.430 | **<.001** |
| *Port*×*Treatment* | 1,48 | 0.096 | .758 | - | - | - |
| *Port*×*Phenotype* | 2,48 | 8.537 | **<.001** | - | - | - |
| *Treatment*×*Phenotype* | 2,48 | 0.365 | .696 | 1,48 | 1.055 | .356 |
| *Port*×*Treatment*×*Phenotype* | 2,48 | 0.623 | .541 | - | - | - |

**Extended Data Table 4.**
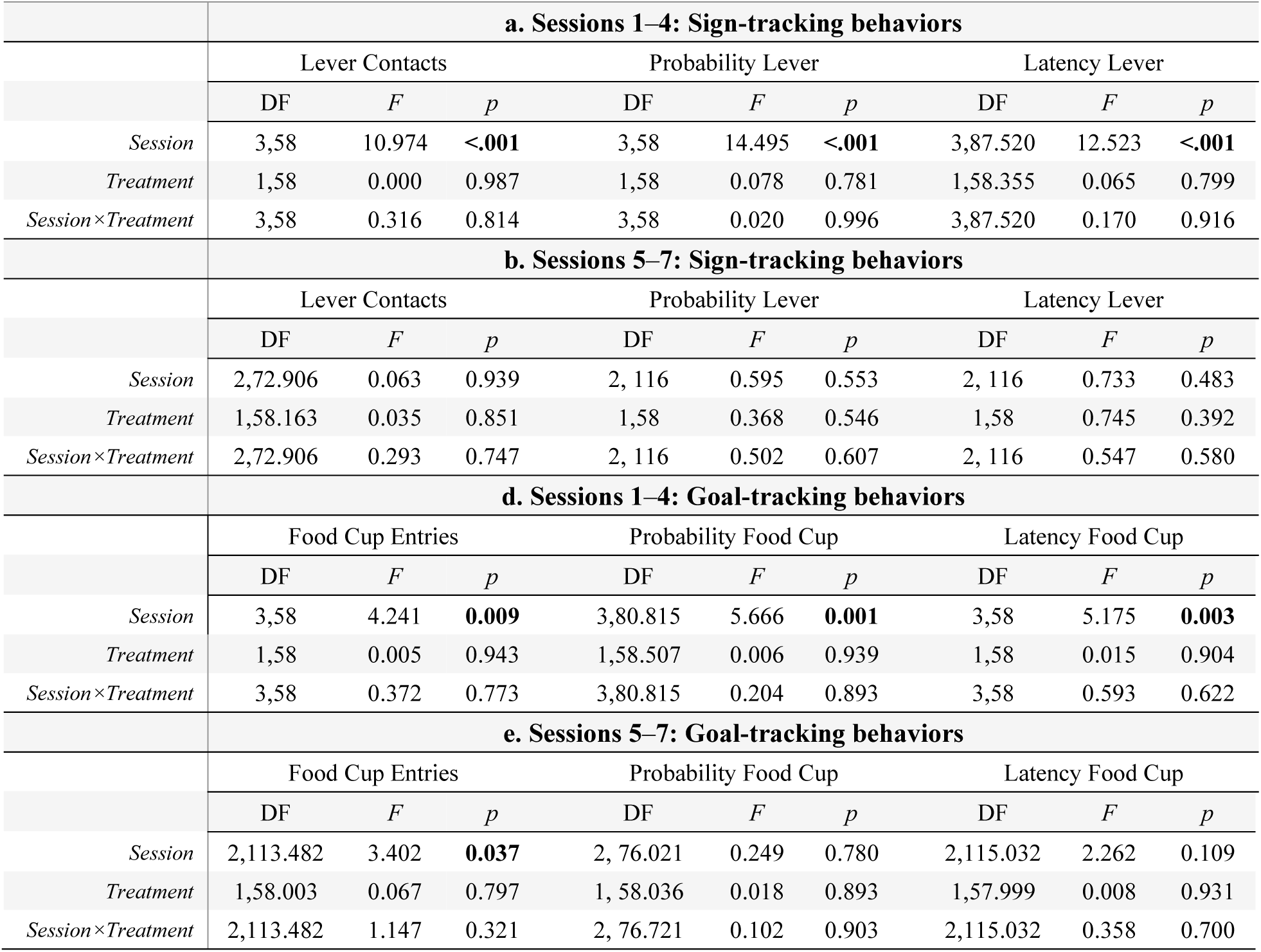
Statistical analyses for sign- and goal-tracking behaviors in the absence of DREADD. Data from linear mixed effects model analyses assessing the effect of treatment in rats that did not have surgery for DREADD expression. Results are shown for (a,d) sessions 1–4 and (b,e) sessions 5–7. Contacts/entries, probability, and latency are represented for (a–b) sign-tracking behaviors and (d–e) goal-tracking behaviors. Significant effects and interactions are bolded. These data correspond to that shown in Extended Data Figure 4.

|  | **a. Sessions 1–4: Sign-tracking behaviors** |  |  |  |  |  |  |  |  |
| --- | --- | --- | --- | --- | --- | --- | --- | --- | --- |
|  | Lever Contacts |  |  | Probability Lever |  |  | Latency Lever |  |  |
|  | DF | *F* | *p* | DF | *F* | *p* | DF | *F* | *p* |
| *Session* | 3,58 | 10.974 | **<.001** | 3,58 | 14.495 | **<.001** | 3,87.520 | 12.523 | **<.001** |
| *Treatment* | 1,58 | 0.000 | 0.987 | 1,58 | 0.078 | 0.781 | 1,58.355 | 0.065 | 0.799 |
| *Session*×*Treatment* | 3,58 | 0.316 | 0.814 | 3,58 | 0.020 | 0.996 | 3,87.520 | 0.170 | 0.916 |
|  | **b. Sessions 5–7: Sign-tracking behaviors** |  |  |  |  |  |  |  |  |
|  | Lever Contacts |  |  | Probability Lever |  |  | Latency Lever |  |  |
|  | DF | *F* | *p* | DF | *F* | *p* | DF | *F* | *p* |
| *Session* | 2,72.906 | 0.063 | 0.939 | 2, 116 | 0.595 | 0.553 | 2, 116 | 0.733 | 0.483 |
| *Treatment* | 1,58.163 | 0.035 | 0.851 | 1,58 | 0.368 | 0.546 | 1,58 | 0.745 | 0.392 |
| *Session*×*Treatment* | 2,72.906 | 0.293 | 0.747 | 2, 116 | 0.502 | 0.607 | 2, 116 | 0.547 | 0.580 |
|  | **d. Sessions 1–4: Goal-tracking behaviors** |  |  |  |  |  |  |  |  |
|  | Food Cup Entries |  |  | Probability Food Cup |  |  | Latency Food Cup |  |  |
|  | DF | *F* | *p* | DF | *F* | *p* | DF | *F* | *p* |
| *Session* | 3,58 | 4.241 | **0.009** | 3,80.815 | 5.666 | **0.001** | 3,58 | 5.175 | **0.003** |
| *Treatment* | 1,58 | 0.005 | 0.943 | 1,58.507 | 0.006 | 0.939 | 1,58 | 0.015 | 0.904 |
| *Session*×*Treatment* | 3,58 | 0.372 | 0.773 | 3,80.815 | 0.204 | 0.893 | 3,58 | 0.593 | 0.622 |
|  | **e. Sessions 5–7: Goal-tracking behaviors** |  |  |  |  |  |  |  |  |
|  | Food Cup Entries |  |  | Probability Food Cup |  |  | Latency Food Cup |  |  |
|  | DF | *F* | *p* | DF | *F* | *p* | DF | *F* | *p* |
| *Session* | 2,113.482 | 3.402 | **0.037** | 2, 76.021 | 0.249 | 0.780 | 2,115.032 | 2.262 | 0.109 |
| *Treatment* | 1,58.003 | 0.067 | 0.797 | 1, 58.036 | 0.018 | 0.893 | 1,57.999 | 0.008 | 0.931 |
| *Session*×*Treatment* | 2,113.482 | 1.147 | 0.321 | 2, 76.721 | 0.102 | 0.903 | 2,115.032 | 0.358 | 0.700 |

**Extended Data Table 5.** Statistical analyses for the effects of treatment and phenotype on sign-tracking behaviors in the absence of DREADD. Data from linear mixed effects model analyses assessing the effect of treatment in rats that did not have surgery for DREADD expression. Results are shown for (a) sessions 1–4 and (b) sessions 5–7, comparing rats by treatment and phenotype. Contacts, probability, and latency are represented for sign-tracking behaviors. Significant effects and interactions are bolded. These data correspond to that shown in Extended Data Figure 4.

|  | **a. Sessions 1–4: Sign-tracking behaviors** |  |  |  |  |  |  |  |  |
| --- | --- | --- | --- | --- | --- | --- | --- | --- | --- |
|  | Lever Contacts |  |  | Probability Lever |  |  | Latency Lever |  |  |
|  | DF | F | p | DF | F | p | DF | F | p |
| *Session* | 3,54 | 19.784 | **<.001** | 3,144.296 | 18.010 | **<.001** | 3,95.656 | 23.674 | **<.001** |
| *Treatment* | 1,54 | 0.019 | 0.890 | 1,55.497 | 0.098 | 0.756 | 1,58.901 | 0.045 | 0.832 |
| *Phenotype* | 2,54 | 34.867 | **<.001** | 2,55.497 | 62.546 | **<.001** | 2,58.901 | 52.097 | **<.001** |
| *Session*×*Treatment* | 3,54 | 1.061 | 0.373 | 3,144.296 | 0.069 | 0.976 | 3,95.656 | 0.208 | 0.891 |
| *Session*×*Phenotype* | 6,54 | 16.765 | **<.001** | 61,44.296 | 10.977 | **<.001** | 6,95.656 | 15.878 | **<.001** |
| *Treatment*×*Phenotype* | 2,54 | 0.178 | 0.838 | 2,55.497 | 0.368 | 0.694 | 2,58.901 | 0.257 | 0.774 |
| *Session*×*Treatment*×*Phenotype* | 6,54 | 0.625 | 0.710 | 6,144.296 | 0.251 | 0.958 | 6,95.656 | 0.227 | 0.967 |
|  | **b. Sessions 5–7: Sign-tracking behaviors** |  |  |  |  |  |  |  |  |
|  | Lever Contacts |  |  | Probability Lever |  |  | Latency Lever |  |  |
|  | DF | F | p | DF | F | p | DF | F | p |
| *Session* | 2,108.463 | 0.003 | 0.997 | 2,108 | 0.404 | .669 | 2,108 | 0.713 | .493 |
| *Treatment* | 1,56.702 | 0.162 | 0.689 | 1,54 | 0.865 | .356 | 1,54 | 1.449 | .234 |
| *Phenotype* | 2,56.702 | 35.261 | **<.001** | 2,54 | 70.615 | **<.001** | 2,54 | 51.708 | **<.001** |
| *Session*×*Treatment* | 2,108.463 | 0.207 | 0.813 | 2,108 | 0.555 | .575 | 2,108 | 0.534 | .588 |
| *Session*×*Phenotype* | 4,108.463 | 2.672 | **0.036** | 4,108 | 2.422 | .053 | 4,108 | 2.100 | .086 |
| *Treatment*×*Phenotype* | 2,56.702 | 1.076 | 0.348 | 2,54 | 0.349 | .707 | 2,54 | 0.079 | .925 |
| *Session*×*Treatment*×*Phenotype* | 4,108.463 | 0.615 | 0.652 | 4,108 | 0.589 | .671 | 4,108 | 1.599 | .180 |

**Extended Data Table 6.**
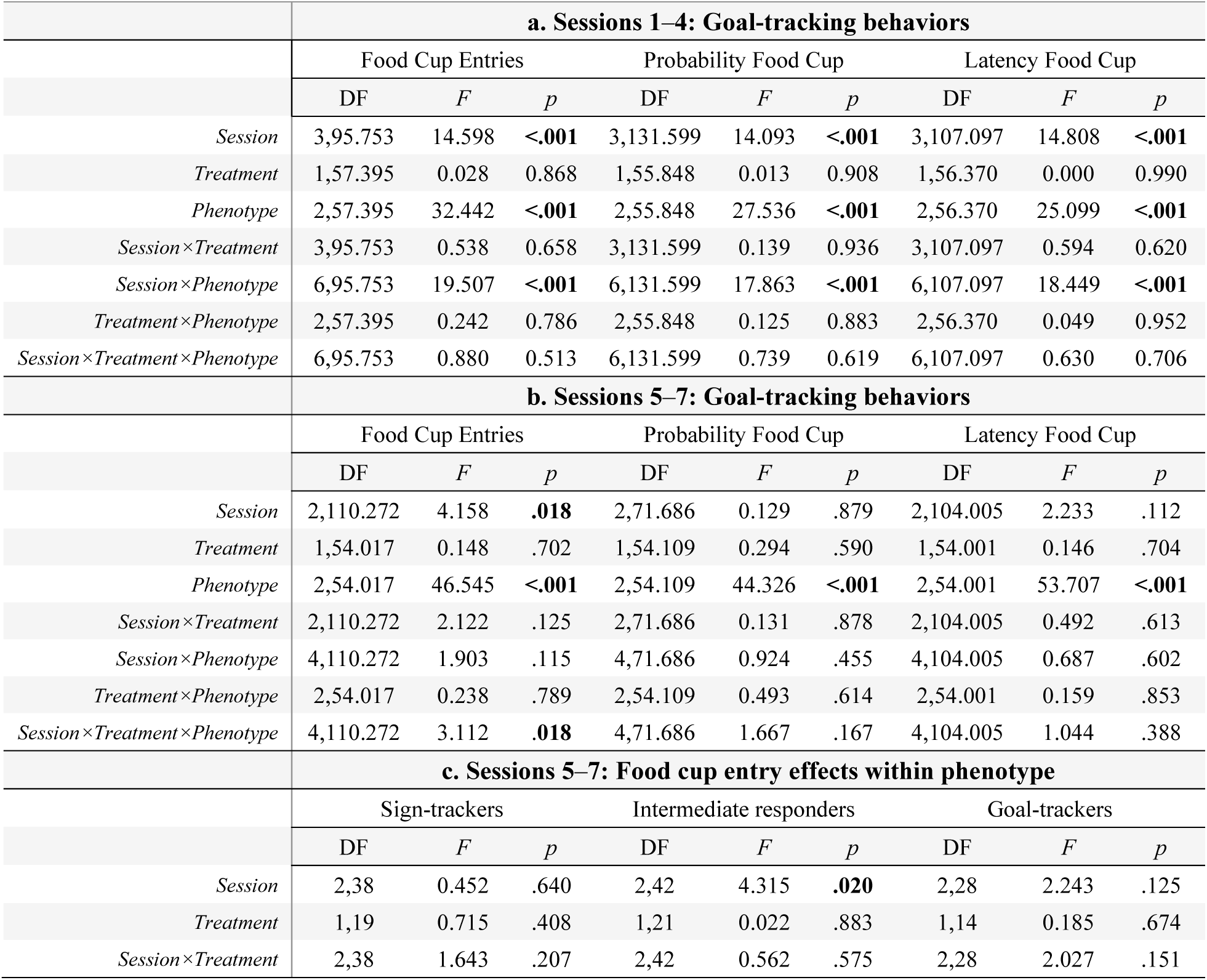
Statistical analyses for the effects of treatment and phenotype on goal-tracking behaviors in the absence of DREADD. Data from linear mixed effects model analyses in rats that did not have surgery for DREADD expression. Data are shown for (a) sessions 1–4 and (b) sessions 5–7, comparing rats by treatment and phenotype, and (c) shows follow-up analyses within each phenotype for the significant session × treatment × phenotype interaction for food cup entries. Entries, probability, and latency are represented for goal-tracking behaviors. Significant effects and interactions are bolded. These data correspond to that shown in Extended Data Figure 4.

|  | **a. Sessions 1–4: Goal-tracking behaviors** |  |  |  |  |  |  |  |  |
| --- | --- | --- | --- | --- | --- | --- | --- | --- | --- |
|  | Food Cup Entries |  |  | Probability Food Cup |  |  | Latency Food Cup |  |  |
|  | DF | *F* | *p* | DF | *F* | *p* | DF | *F* | *p* |
| *Session* | 3,95.753 | 14.598 | **<.001** | 3,131.599 | 14.093 | **<.001** | 3,107.097 | 14.808 | **<.001** |
| *Treatment* | 1,57.395 | 0.028 | 0.868 | 1,55.848 | 0.013 | 0.908 | 1,56.370 | 0.000 | 0.990 |
| *Phenotype* | 2,57.395 | 32.442 | **<.001** | 2,55.848 | 27.536 | **<.001** | 2,56.370 | 25.099 | **<.001** |
| *Session* × *Treatment* | 3,95.753 | 0.538 | 0.658 | 3,131.599 | 0.139 | 0.936 | 3,107.097 | 0.594 | 0.620 |
| *Session* × *Phenotype* | 6,95.753 | 19.507 | **<.001** | 6,131.599 | 17.863 | **<.001** | 6,107.097 | 18.449 | **<.001** |
| *Treatment* × *Phenotype* | 2,57.395 | 0.242 | 0.786 | 2,55.848 | 0.125 | 0.883 | 2,56.370 | 0.049 | 0.952 |
| *Session* × *Treatment* × *Phenotype* | 6,95.753 | 0.880 | 0.513 | 6,131.599 | 0.739 | 0.619 | 6,107.097 | 0.630 | 0.706 |
|  | **b. Sessions 5–7: Goal-tracking behaviors** |  |  |  |  |  |  |  |  |
|  | Food Cup Entries |  |  | Probability Food Cup |  |  | Latency Food Cup |  |  |
|  | DF | *F* | *p* | DF | *F* | *p* | DF | *F* | *p* |
| *Session* | 2,110.272 | 4.158 | **.018** | 2,71.686 | 0.129 | .879 | 2,104.005 | 2.233 | .112 |
| *Treatment* | 1,54.017 | 0.148 | .702 | 1,54.109 | 0.294 | .590 | 1,54.001 | 0.146 | .704 |
| *Phenotype* | 2,54.017 | 46.545 | **<.001** | 2,54.109 | 44.326 | **<.001** | 2,54.001 | 53.707 | **<.001** |
| *Session* × *Treatment* | 2,110.272 | 2.122 | .125 | 2,71.686 | 0.131 | .878 | 2,104.005 | 0.492 | .613 |
| *Session* × *Phenotype* | 4,110.272 | 1.903 | .115 | 4,71.686 | 0.924 | .455 | 4,104.005 | 0.687 | .602 |
| *Treatment* × *Phenotype* | 2,54.017 | 0.238 | .789 | 2,54.109 | 0.493 | .614 | 2,54.001 | 0.159 | .853 |
| *Session* × *Treatment* × *Phenotype* | 4,110.272 | 3.112 | **.018** | 4,71.686 | 1.667 | .167 | 4,104.005 | 1.044 | .388 |
|  | **c. Sessions 5–7: Food cup entry effects within phenotype** |  |  |  |  |  |  |  |  |
|  | Sign-trackers |  |  | Intermediate responders |  |  | Goal-trackers |  |  |
|  | DF | *F* | *p* | DF | *F* | *p* | DF | *F* | *p* |
| *Session* | 2,38 | 0.452 | .640 | 2,42 | 4.315 | **.020** | 2,28 | 2.243 | .125 |
| *Treatment* | 1,19 | 0.715 | .408 | 1,21 | 0.022 | .883 | 1,14 | 0.185 | .674 |
| *Session* × *Treatment* | 2,38 | 1.643 | .207 | 2,42 | 0.562 | .575 | 2,28 | 2.027 | .151 |

**Extended Data Table 7.** Statistical analyses for behavior during the Conditioned Reinforcement Test (CRT) in the absence of DREADD. Data from a mixed ANOVA comparing (left column) nose pokes into each port and a two-way ANOVA comparing lever contacts during the conditioned reinforcement test. Analysis was conducted between (a) treatment groups and (b) with treatment and phenotype as variables. Significant effects and interactions are bolded. These data correspond to that shown in Extended Data Figure 5.

|  | **a. Session 8: Conditioned Reinforcement** |  |  |  |  |  |
| --- | --- | --- | --- | --- | --- | --- |
|  | Nose Pokes |  |  | Lever Contacts |  |  |
|  | DF | *F* | *p* | DF | *F* | *p* |
| *Port* | 1,58 | 22.034 | **<.001** | - | - | - |
| *Treatment* | 1,58 | 0.538 | .466 | 1,58 | 0.832 | .365 |
| *Port*×*Treatment* | 1,58 | 0.163 | .687 | - | - | - |
|  | **b. Session 8: Conditioned Reinforcement** |  |  |  |  |  |
|  | Nose Pokes |  |  | Lever Contacts |  |  |
|  | DF | *F* | *p* | DF | *F* | *p* |
| *Port* | 1,54 | 21.243 | **<.001** | - | - | - |
| *Treatment* | 1,54 | 0.990 | .324 | 1,54 | 0.369 | .546 |
| *Phenotype* | 2,54 | 3.648 | **.033** | 2,54 | 10.304 | **<.001** |
| *Port*×*Treatment* | 1,54 | 0.095 | .759 | - | - | - |
| *Port*×*Phenotype* | 2,54 | 2.848 | .067 | - | - | - |
| *Treatment*×*Phenotype* | 2,54 | 0.677 | .512 | 1,54 | 0.062 | .940 |
| *Port*×*Treatment*×*Phenotype* | 2,54 | 0.255 | .776 | - | - | - |

